# Extensive characterization of HIV-1 reservoirs reveals links to plasma viremia before and during analytical treatment interruption

**DOI:** 10.1101/2021.02.04.429690

**Authors:** Basiel Cole, Laurens Lambrechts, Zoe Boyer, Ytse Noppe, Marie-Angélique De Scheerder, John-Sebastian Eden, Bram Vrancken, Timothy E. Schlub, Sherry McLaughlin, Lisa M. Frenkel, Sarah Palmer, Linos Vandekerckhove

## Abstract

The HIV-1 reservoir is composed of cells harboring latent proviruses that are capable of contributing to viremia upon antiretroviral treatment (**ART)** interruption. Although this reservoir is known to be maintained by clonal expansion, the contribution of large, infected cell clones to residual viremia and viral rebound remains underexplored. Here, we conducted an extensive analysis on four ART-treated individuals who underwent an analytical treatment interruption (**ATI**). We performed subgenomic (V1-V3 *env*), near full-length proviral and integration site sequencing, and used multiple displacement amplification to sequence both the integration site and provirus from single HIV-infected cells. We found eight proviruses that could phylogenetically be linked to plasma virus obtained before or during the ATI. This study highlights a role for HIV-infected cell clones in the maintenance of the replication-competent reservoir and suggests that infected cell clones can directly contribute to rebound viremia upon ATI.

## Introduction

HIV-1 infection remains incurable due to the presence of a persistent viral reservoir, capable of contributing to viral rebound upon treatment interruption (**TI**) (Chun and Fauci, 1999; Chun et al., 1997, 1998; Finzi et al., 1997). Despite efforts to better understand the dynamics and persistence of the HIV-1 viral reservoir, pinpointing the origins of rebounding viruses remains elusive (De Scheerder et al., 2019). Previously, it was shown that infected CD4 T cells can undergo clonal expansion, contributing to the long-term persistence of the HIV-1 viral reservoir during antiretroviral therapy (**ART**) (Boritz et al., 2016; Cohn et al., 2015a; Einkauf et al., 2018; Hosmane et al., 2017; Maldarelli et al., 2014; Salantes et al., 2018; Simonetti et al., 2016; Wagner et al., 2014a; Wang et al., 2018). The observation that low-level viremia (**LLV**) under ART (Aamer et al., 2020; Bailey et al., 2006; Brennan et al., 2009; Halvas et al., 2020; Tobin et al., 2005; Wagner et al., 2013) and rebound viremia upon TI (Aamer et al., 2020; Kearney et al., 2016; Lu et al., 2018a; De Scheerder et al., 2019) often consist of monotypic populations of viruses, suggest that HIV-1 infected cell clones are key contributors to refueling viremia during TI. Clonality of infected cells has historically been demonstrated by recovering identical proviral sequences or identical integration sites (IS) in multiple cells (Cohn et al., 2015b; Hiener et al., 2017; Lee et al., 2017; Maldarelli et al., 2014; Pinzone et al., 2019; Stockenstrom et al., 2015; Wagner et al., 2014b). While the former method allows for qualitative assessment of the proviral genome, it is often not adequate to confidently predict clonal expansion of HIV-1 infected cells, especially when evaluating a short subgenomic region (Lambrechts et al., 2020; Laskey et al., 2016). On the other hand, analysis of IS provides direct proof of clonal expansion, though it typically leaves the proviral sequence uncharacterized. Recently, three techniques to link near full-length (**NFL**) proviral sequences to IS were developed by Einkauf *et al*. (Einkauf et al., 2018), Patro *et al*. (Patro et al., 2019) and Artesi *et al*. (Artesi et al., 2021), respectively called Matched Integration site and Proviral sequencing (**MIP-Seq**), Multiple Displacement Amplification Single-Genome Sequencing (**MDA-SGS**) and Pooled CRISPR Inverse PCR sequencing (**PCIP-seq**). These assays combine the qualitative strength of NFL HIV-1 sequencing with IS analysis, shedding light on the integration profile of intact versus defective proviruses.

Analytical treatment interruption (**ATI**) studies allow for the investigation of the dynamics and genetic makeup of rebounding viruses (Clarridge et al., 2018; Garner et al., 2017; Kearney et al., 2016; Pannus et al., 2020). To identify the source of rebounding viruses, we previously conducted the HIV-STAR (HIV-1 sequencing before analytical treatment interruption to identify the anatomically relevant HIV reservoir) study (De Scheerder et al., 2019). During this study, in-depth sampling was performed on 11 chronically treated HIV-1 infected participants prior to ATI. Cells were isolated from different anatomical compartments and sorted into several CD4 T cell subsets. Subgenomic proviral sequences (V1-V3 region of *env*) were recovered and phylogenetically linked to sequences from rebounding plasma virus collected during different stages of the ATI. This study suggested that HIV-1 rebound is predominantly fueled by genetically identical viral expansions, highlighting the potentially important role of clonal expansion in the maintenance of the HIV-1 reservoir. While this study yielded a total of 4329 V1-V3 *env* sequences from peripheral blood mononuclear cells (**PBMCs**), lymph node (**LN**) and gut-associated lymphoid tissue **(GALT**), enabling a detailed investigation of the viral reservoir and its relation to rebound viremia, it left some questions unanswered. Most importantly, the evaluation of a short subgenomic region (V1-V3 *env*) to link proviral sequences to rebounding plasma virus made it impossible to investigate the entire genome structure of proviruses linked to rebound. Furthermore, the lack of integration site (IS) analysis did not allow for the study of the chromosomal location of the rebounding proviruses.

To address these points, we performed a combination of multiple displacement amplification (**MDA**), IS analysis and NFL proviral sequencing on four participants that were enrolled in the HIV-STAR study, with special attention to clonally expanded HIV-1 infected cells. We demonstrate that HIV-1 proviral sequences and corresponding IS of clonally expanded infected cells could be retrieved, and in rare cases these could be linked to low-level viremia during ART and rebound viremia upon ATI, highlighting the clinical relevance of large, infected cell clones.

## Results

### Experimental set-up

To investigate the genetic composition and chromosomal location of proviruses within clonally expanded cells, and their relationship to rebound viremia, several qualitative assays were performed on samples from chronically treated HIV-1 infected individuals undergoing an ATI (Figure 1A, Table S1). Samples from these individuals were obtained longitudinally before (T1) and during the ATI (T2, T3, T4), as summarized in Figure 1B.

**Figure 1:**
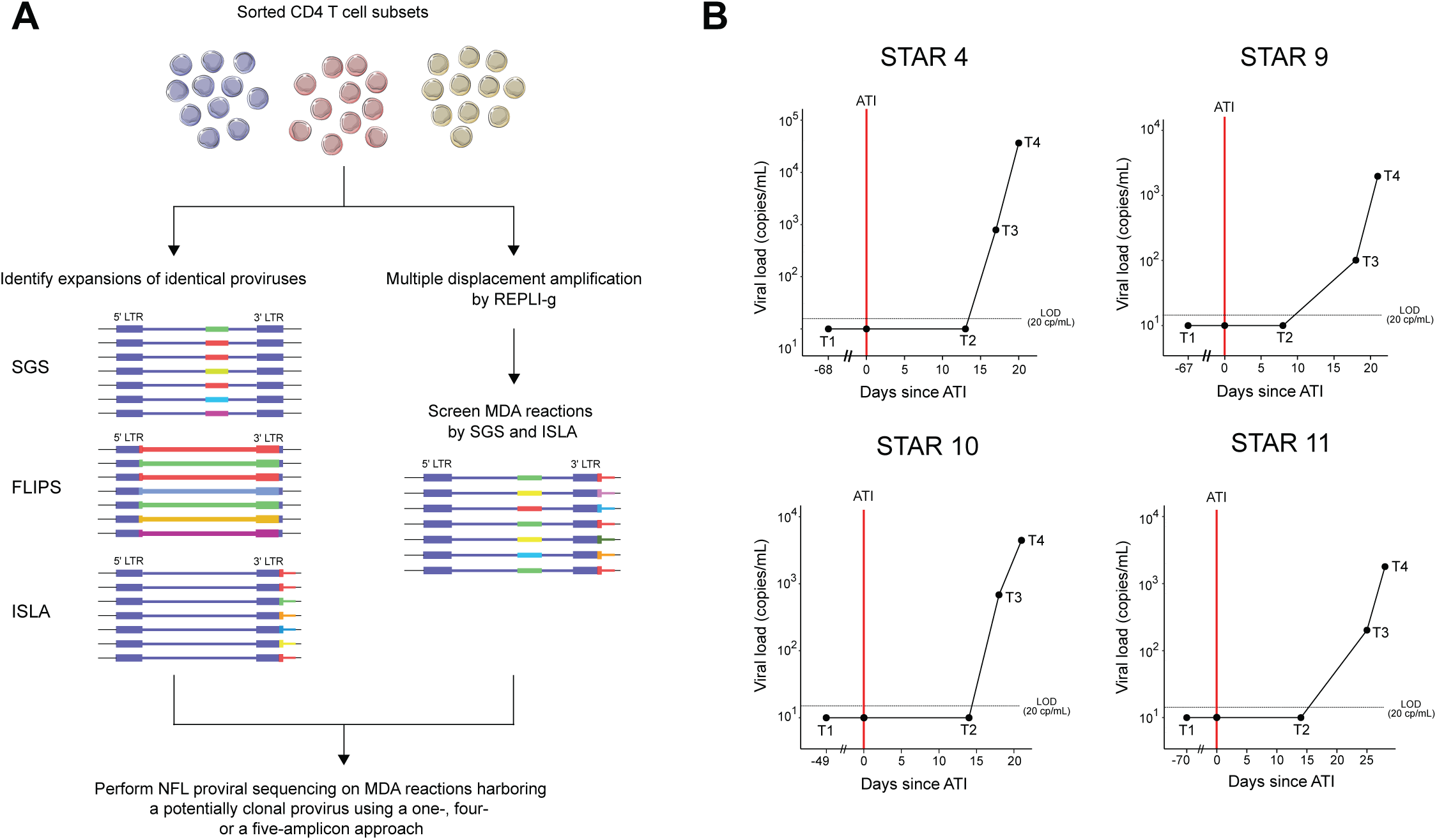
Overview of the workflow for HIV-1 reservoir characterization and viral loads at each timepoint of sample collection. (A) Workflow of HIV-1 reservoir characterization by single-genome sequencing (**SGS**), full-length individual proviral sequencing (**FLIPS**), integration site loop amplification (**ISLA**) and multiple displacement amplification (**MDA**). In a first step, potentially clonal HIV-1 infected cells were identified by SGS, FLIPS and ISLA on lysed sorted CD4 T-cell subsets. In a second step, MDA with subsequent SGS and ISLA was performed on selected sorted cell lysates. In the final step, MDA reactions containing a potentially clonal provirus were identified and the near full-length (**NFL**) genome of the according provirus was amplified and sequenced. (B) Viral load (copies/mL) at each time of sample collection for all participants. The day of analytical treatment interruption (**ATI**) initiation is indicated with a vertical red line. The plasma was sampled during ART (timepoint 1, T1), 8 to 14 days after ATI (timepoint 2, T2), at the first detectable viral load (timepoint 3, T3), and at later rebound (timepoint 4, T4). Note that T1 is not shown to scale. The horizontal dashed lines indicate the limit of detection at 20 copies/mL.

First, the overall landscape of HIV-1 infected cell clones prior to ATI (T1, Figure 1B) was determined by subgenomic single-genome sequencing (**SGS**) and Full-length Individual Proviral Sequencing (**FLIPS**) of proviruses, and by Integration Site Loop Amplification (**ISLA**) at the integration site level (Figure 1A, Table S1). This yielded three datasets that were used independently to identify potential clonally expanded infected cell populations.

Next, to find links between the different datasets, MDA was performed at limiting dilution on sorted cell lysates from peripheral blood obtained before ATI (T1). MDA wells were subjected to V1-V3 *env* SGS and ISLA, and MDA reactions that yielded a V1-V3 *env* sequence and/or an IS corresponding to a suspected cellular clone were further investigated. This was determined by an exact link to ISLA/FLIPS/SGS data generated in the first step, or by identical V1-V3 *env* sequences and/or IS shared between several MDA wells. The NFL genomes of the proviruses in these selected MDA wells were amplified and sequenced using either a one-amplicon, four-amplicon, or five-amplicon approach, or a combination thereof (see Methods). These MDA-derived NFL sequences were subsequently mapped back to proviral FLIPS sequences and historic V1-V3 *env* proviral sequences from PBMCs, GALT and LN subsets prior to ATI (T1, Figure 1B), as well as V1-V3 *env* plasma-derived RNA sequences retrieved during the ATI (T2-T4, Figure 1B). In addition, remaining T4 plasma samples from all four participants were subjected to 5’- and 3’-half genome amplification to complement existing V1-V3 *env* plasma SGS data.

This set-up allowed for the assessment of the genetic structure of proviruses in clonally expanded infected cells, their placement across cellular subsets and anatomical compartments, and their contribution to residual viremia on ART and rebound viremia during an ATI.

### Integration site analysis and full-length proviral sequencing

To gain insight into the composition of the viral reservoirs of the four STAR participants, especially in terms of clonal expansion of infected cells, we initially performed bulk NFL proviral sequencing and IS analysis.

ISLA was performed on endpoint-diluted non-amplified cell lysate and on MDA-amplified cell lysates of central memory/transitional memory (**TCM/TTM**) and effector memory (**TEM**) CD4 T cell subsets from peripheral blood (T1) from three of the four participants: STAR 9, STAR 10 and STAR 11 (Figure 2A, Tables S2 and S3). Analysis of IS revealed a significantly higher degree of clonally expanded HIV-1 infected cells in the TEM subset (mean 55%) compared to the TCM/TTM fraction (mean 16%) in the peripheral blood (P < 0.001 for STAR 9 and STAR 11; P < 0.05 for STAR 10), as previously reported (Hiener et al., 2017). Identical IS between different subsets, suggestive of linear differentiation from an originally infected TCM/TTM into a TEM, was observed in rare instances, with IS at 8 unique positions in the human genome shared between subsets out of 328 IS recovered (171 in TCM/TTM and 157 in TEM).

**Figure 2:**
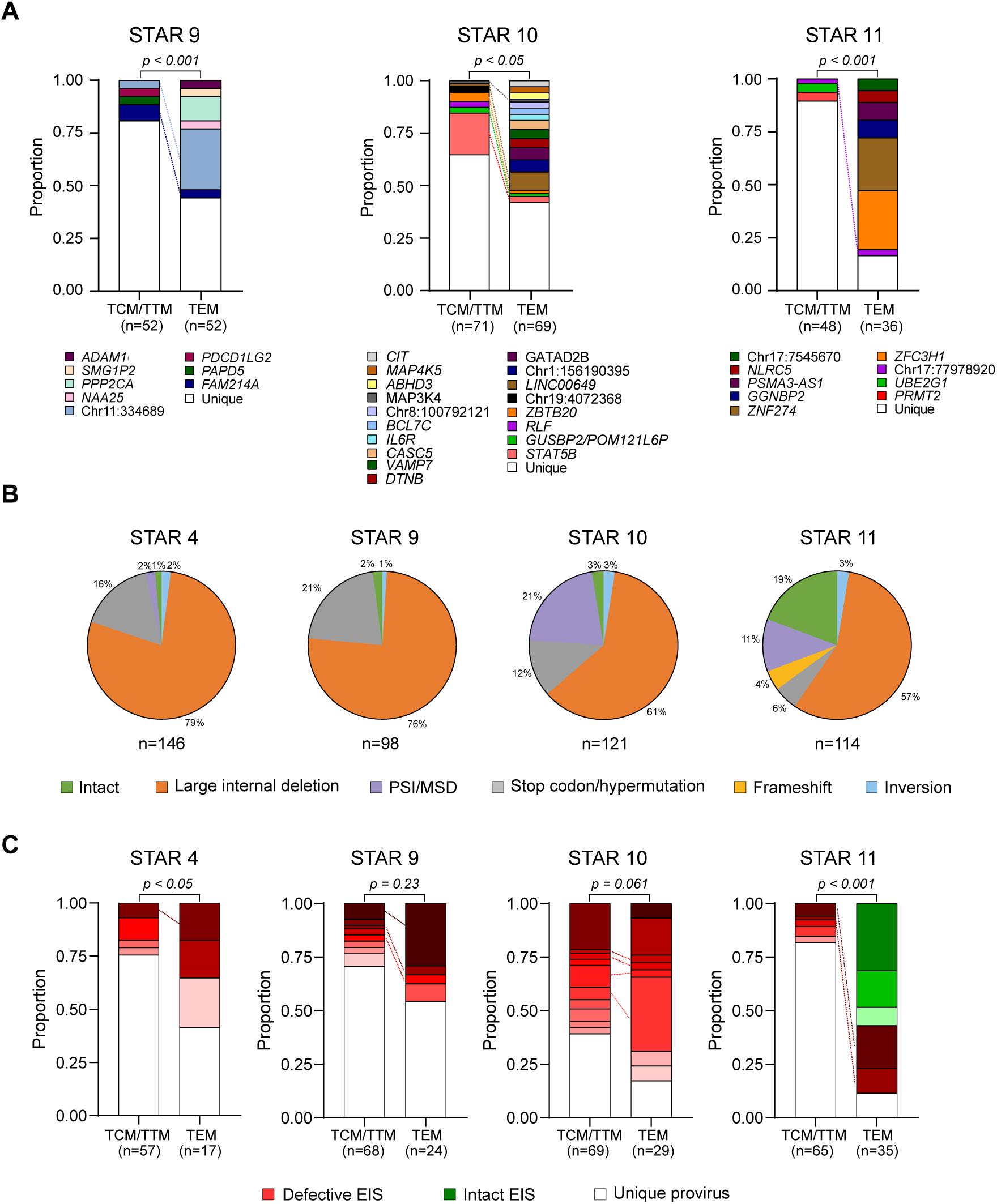
Classification of HIV-1 integration sites and proviral near full-length genome sequences across different cell subsets before ATI. (A) Proportions of integration sites (**IS**) retrieved by integration site loop amplification (**ISLA**) for participants STAR 9, STAR 10 and STAR 11 from TCM/TTM and TEM subsets from peripheral blood. IS found more than once are defined as “clonal” and are shown in color as proportion of all IS. Identical IS found in both subsets are linked with dashed lines. P-values test was used for a difference in the proportion of unique IS between TCM/TTM and TEM by “prop.test” in R. (B) Proportions of intact and defective near full-length sequences from Full-Length Individual Provirus sequencing (**FLIPS**) within all sequenced proviruses from peripheral blood, gut-associated lymphoid tissue (**GALT)** and lymph nodes for each participant. (C) Proportions of identical sequences (**EIS**) found in TCM/TTM and TEM peripheral blood subsets based on FLIPS data for each participant. EIS consisting of a detective and intact provirus are shown in shades of red and green respectively while unique proviruses are grouped in the unique provirus category. P-values test was used for a difference in the proportion of unique proviruses between TCM/TTM and TEM by “prop.test” in R. TCM/TTM = central/transitional memory CD4 T cell, TEM = effector memory CD4 T cell.

Near full-length provial sequencing (spanning 92% of the proviral genome) was performed on the TCM/TTM and TEM subsets in the peripheral blood and on CD45+ cells from the GALT for all four participants (T1, Figure 1B). In addition, based on sample availability, other cell subsets from the peripheral blood and LN were assayed with FLIPS for some participants, as listed in Table S1. This yielded a total number of 479 proviral genomes with a mean of 120 genomes per participant (Figure 2B, Table S3). Across all participants, 29 (6%) intact proviral genomes were retrieved, with a majority of proviral sequences (68%, n=327) displaying large internal deletions (Figures S1 and S2A). Overall, intact proviral genomes were found more frequently in the TEM fraction compared to the TCM/TTM fraction (Figure S2D). In addition, the overall HIV-1 infection frequency differed significantly between lymphocyte cell subsets from the peripheral blood (P < 0.001), with the TEM subset having the highest infection frequency, except for participant STAR 4 (Figure S2C). Expansions of identical sequences (EIS), suggestive of clonal expansion of infected cells, were observed in all participants (Figure 2C). In accordance with the ISLA results, EIS were more frequent in the TEM fraction compared to the TCM/TTM fraction (P < 0.05 for STAR 4; P = 0.23 for STAR 9; P = 0.061 for STAR 10; P < 0.001 for STAR 11, Figure 2C).

### Multiple displacement amplification-mediated characterization of near full-length proviruses

MDA-mediated HIV-1 provirus and IS sequencing offers the unique opportunity of linking NFL proviral sequences to their precise location in a chromosome. Applying this technique to three of the four study participants, we detected several expanded clones from which NFL sequences and linked IS were sequenced (n=12), as shown in Figure 3A (Table S3).

**Figure 3:**
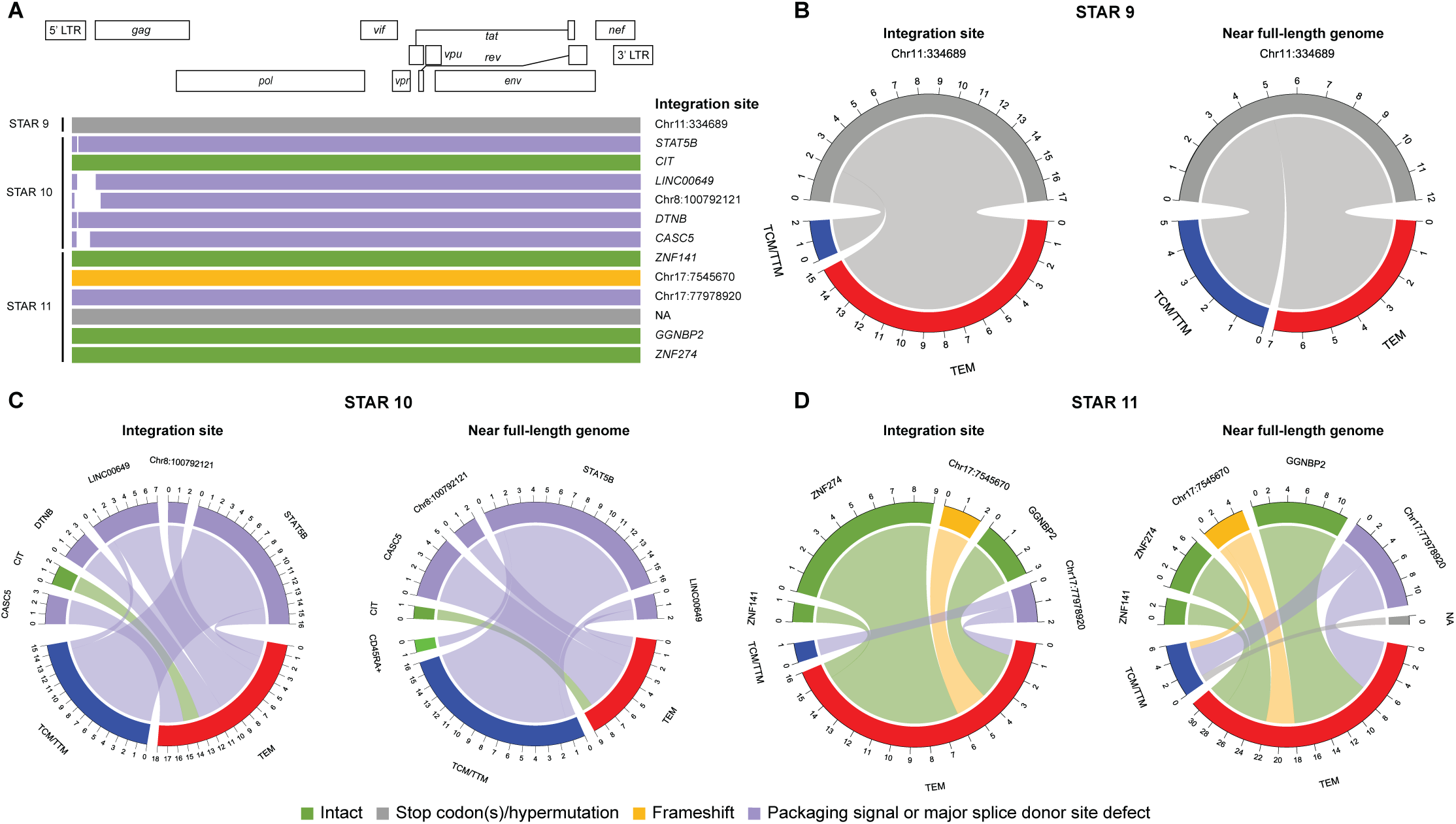
Near full-length proviral HIV-1 genomes and associated integration sites recovered from the peripheral blood by MDA. (A) For each participant, the recovered proviral genome structures are shown aligned to the HXB2 reference genome and corresponding integration sites, if available, are listed on the right. Each provirus is colored according to their structural category. (B-D) For each participant, the number of integration site (**IS**) and near full-length (**NFL**) proviruses linked to each multiple displacement amplification (**MDA**) clone are shown together with their corresponding cellular subset. TCM/TTM = central/transitional memory CD4 T cell, TEM = effector memory CD4 T cell, NA = not available

In total, 8 clonal cell populations with a defective provirus were identified: 1 with hypermutation, 1 with a frameshift, and 6 with a defect in the packaging signal and/or major splice donor site (Figure 3A). Somewhat less prevalent were clones with a putatively intact proviral genome (n=4), 3 of these were retrieved in STAR 11 (Figure 3A). Interestingly, 5 out of 8 defective clones were found in both the TCM/TTM and the TEM fractions, which suggests linear differentiation of clonal cell populations (Figures 3B-3D). In contrast, all the clones with intact proviruses were found exclusively in the TEM fraction (Figures 3C and 3D).

Of note, we identified one cell clone with an IS in *STAT5B*, harboring a provirus with a 25-bp deletion in the packaging signal (Figure 3A). Previous work has shown that proviral integration into the first intron of this gene in the *cis* configuration can lead to HIV LTR-driven dysregulation of *STAT5B*, resulting in cellular proliferation (Cesana et al., 2017). However, in the present study, the provirus was integrated in the first intron in the *trans* configuration, suggesting that integration in the *trans* configuration can also result in aberrant transcription of *STAT5B*, or that the clone was expanded by mechanisms unrelated to the IS.

We conclude that a large fraction of the clonally expanded infected cell populations we identified harbor defective proviruses and that these are frequently found across different cellular subsets, suggesting differentiation of infected cell populations.

### Large discrepancies between suspected clonal HIV-1 infected cell populations identified with ISLA, SGS and FLIPS

ISLA, SGS and FLIPS can independently be used to assess clonality of infected cells, the former based on the integration site and the two latter on a (subgenomic) sequence of the proviral genome. To investigate whether the methods appear biased in their ability to detect specific clones, we used V1-V3 *env* or NFL sequences to assess overlap between assays (Figure 4). In the case of IS data, only those IS that were associated with a corresponding V1-V3 *env* sequence (identified through MDA) could be linked to other assays. For FLIPS sequences, proviruses that had an internal deletion covering the V1-V3 *env* region could not be linked to SGS data. Matches between IS (upper panel) and FLIPS data (bottom panel) were based on NFL sequences, whereas other links were based on V1-V3 *env* sequences (IS and SGS; SGS and FLIPS). For matches between NFL sequences, up to 3-bp differences were allowed to account for PCR-induced errors and sequencing errors, while 100% concordance was required for V1-V3 *env* matches. To assess the capacity of the V1-V3 *env* region to distinguish different NFL proviruses, the clonal prediction score (**CPS**) was calculated for each participant(Laskey et al., 2016) (Table S5). In addition, the nucleotide diversity of the V1-V3 *env* region was calculated (Table S5).

**Figure 4:**
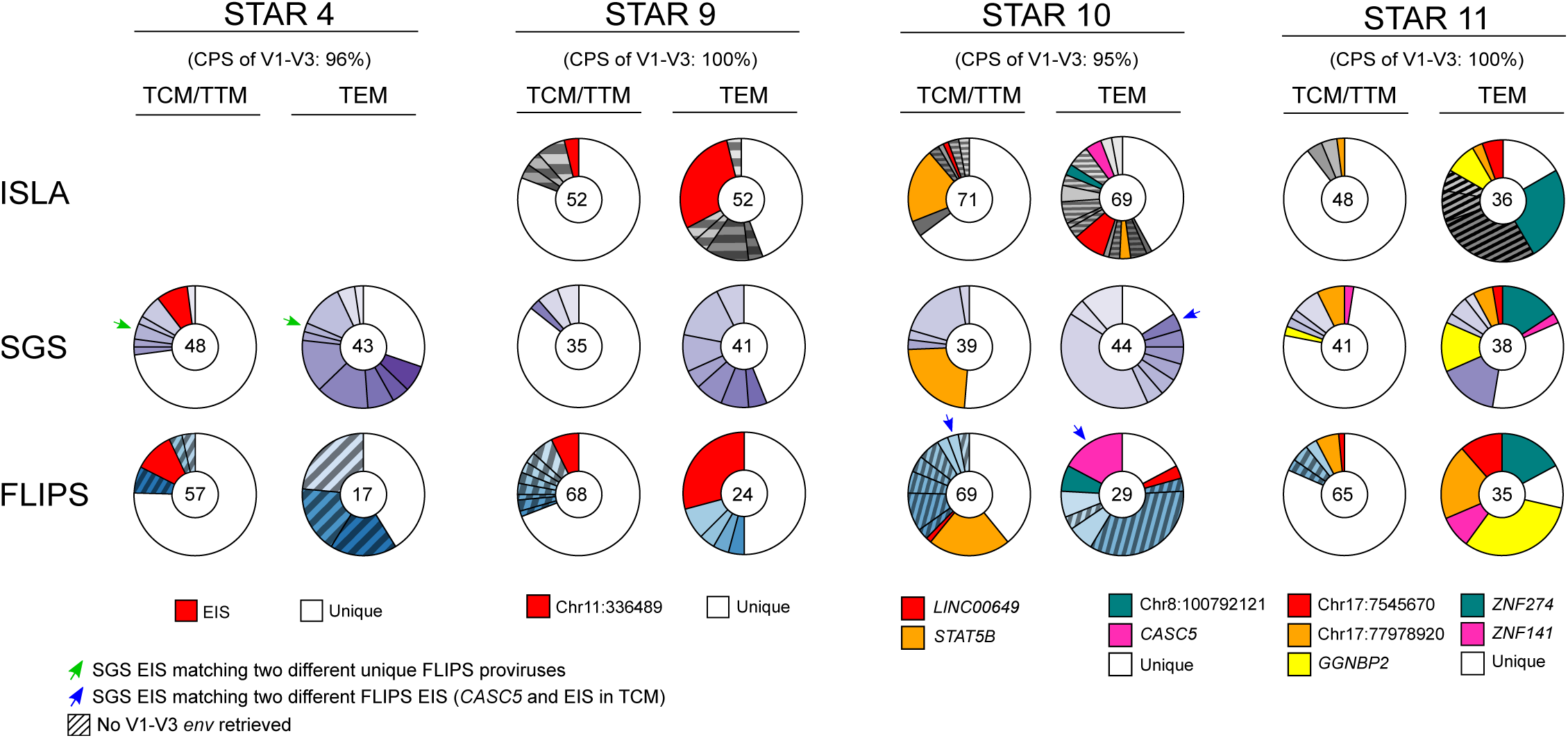
Comparison of assays to identify potentially clonal HIV-1 infected cell populations. The total number of examined integration sites (**IS**), V1-V3 *env* sequences and near full-length (**NFL**) proviral sequences is noted in the middle of each donut plot. Sequences found multiple times within the same assay are colored by a shade of grey, purple or blue (for integration site loop amplification (ISLA), single-genome sequencing (**SGS**) and full-length individual proviral sequencing (**FLIPS**) respectively). When NFL or V1-V3 *env* sequences could be linked to an identified multiple displacement amplification (**MDA**) cell clone, they were given a distinct standout color and chromosome designation as indicated in the legend. Populations of identical FLIPS or ISLA sequences that are not associated with a V1-V3 *env* sequence (due to deletions and/or primer mismatches) are shaded. Arrows are used to indicate discrepancies between the different assays. EIS = expansion of identical sequences.

Upon comparison of EIS present in SGS data and FLIPS data from STAR 4, one clear overlap could be found in the peripheral blood TCM/TTM fraction. All other proviral sequences retrieved with SGS could not be linked unequivocally to EIS identified using FLIPS, indicating a significant bias. However, one V1-V3 *env* sequence found with SGS in the TEM and the TCM/TTM fractions matched two different unique FLIPS sequences (Figure 4, green arrows). This is an example of a presumed clonal EIS as detected by SGS that consists of two or more proviruses sharing the same V1-V3 *env* region, although differing elsewhere in their genome. This discrepancy is reflected by a CPS of 96%, with a V1-V3 *env* nucleotide diversity of 0.01 (Table S5).

A similar picture was observed for STAR 9, with only a single overlap between IS and FLIPS data. One major clone, with a hypermutated provirus integrated at an intergenic region on chromosome 11, was detected with both MDA and FLIPS. In both assays, this clone was predominantly found in the peripheral blood TEM fraction (23% and 29% respectively), but also appeared in the peripheral blood TCM/TTM fraction. However, this provirus was never amplified with SGS, which can be explained by the fact that V1-V3 *env* primers did not anneal to this hypermutated sequence. The CPS for STAR 9 was 100%, with a V1-V3 *env* nucleotide diversity of 0.017 (Table S5).

Participant STAR 10 displayed a clear discrepancy between the assays. One small EIS from the TEM SGS data was shared an identical V1-V3 *env* with two different EIS found in the FLIPS data, one of which can be linked to the *CASC5* cell clone (Figure 4, blue arrows). This is most likely the result of multiple distinct proviruses sharing an identical V1-V3 *env* sequence but integrated at different sites. This resulted in a lower CPS of 95%, with a V1-V3 *env* nucleotide diversity of 0.014 (Table S5).

In contrast with the previous participants, remarkable consistency between assays was observed for STAR 11, with all the clonal NFL sequences identical to those from both SGS and MDA-derived IS data (Figure 4). However, the largest clone based on ISLA data, integrated in the *ZFC3H1* gene (Figure 2A), could not be linked to SGS and FLIPS data, which was probably the result of large internal deletions spanning the entire length of the genome. In fact, out of ten MDA wells that yielded this integration site, the proviral sequence could not be amplified by V1-V3 *env* SGS, or by a one-, four- or five-amplicon approach NFL sequencing (Table S3). The CPS for STAR 11 was 100%, with a V1-V3 *env* nucleotide diversity of 0.017 (Table S5).

In conclusion, we demonstrate that for two out of four participants, the CPS is lower than 100%, suggesting linkage inaccuracies when using the V1-V3 *env* to predict clonality of infected cells. Furthermore, we show compartmentalization between the viral populations identified by the V1-V3 env SGS method versus the FLIPS method. This could either result from primer bias or from sampling bias, caused by the high frequency of env-deleted proviruses sampled by FLIPS.

### Plasma viral sequences match intact proviruses and proviruses with large internal deletions or defects in the packaging signal

In our previous study on HIV-STAR participants, proviral V1-V3 *env* SGS sequences from multiple lymphocyte subsets and anatomical tissues collected prior to ATI were linked to plasma RNA sequences of rebounding viruses (De Scheerder et al., 2019). Yet, no conclusions about the genomic structure of the NFL proviruses and their associated IS could be inferred, since these subgenomic sequences did not allow for such analysis. Data generated in the present study allowed for a deeper characterization of the proviral landscape through linkage of NFL proviral sequences and their corresponding IS to plasma RNA sequences of rebounding viruses.

To investigate how the new FLIPS- and MDA-derived NFL sequences compare to the historic plasma and proviral V1-V3 *env* sequences generated during the original HIV-STAR study(De Scheerder et al., 2019), the trimmed V1-V3 *env* region from the MDA- and FLIPS- derived NFL sequences were aligned with SGS-derived and MDA-derived V1-V3 *env* sequences. Subsequently, phylogenetic trees were constructed for each participant (Figure 5). As seen before in Figure 4, the V1-V3 *env* regions of some NFL proviruses are an identical match to historic proviral V1-V3 *env* SGS sequences and cluster together on the same branch (Figures 5 and S4). Interestingly, identical sequences were detected in different anatomical compartments. For example, participant STAR 9 displays an intact FLIPS provirus which falls into a cluster of proviral peripheral blood and GALT V1-V3 *env* sequences (Figure 5B). Furthermore, participant STAR 10 has an MDA-derived provirus integrated in *STAT5B* that matches the V1-V3 *env* proviral sequences from the peripheral blood and LN, suggesting that cells of this clone traffic between these compartments (Figure 5C).

**Figure 5:**
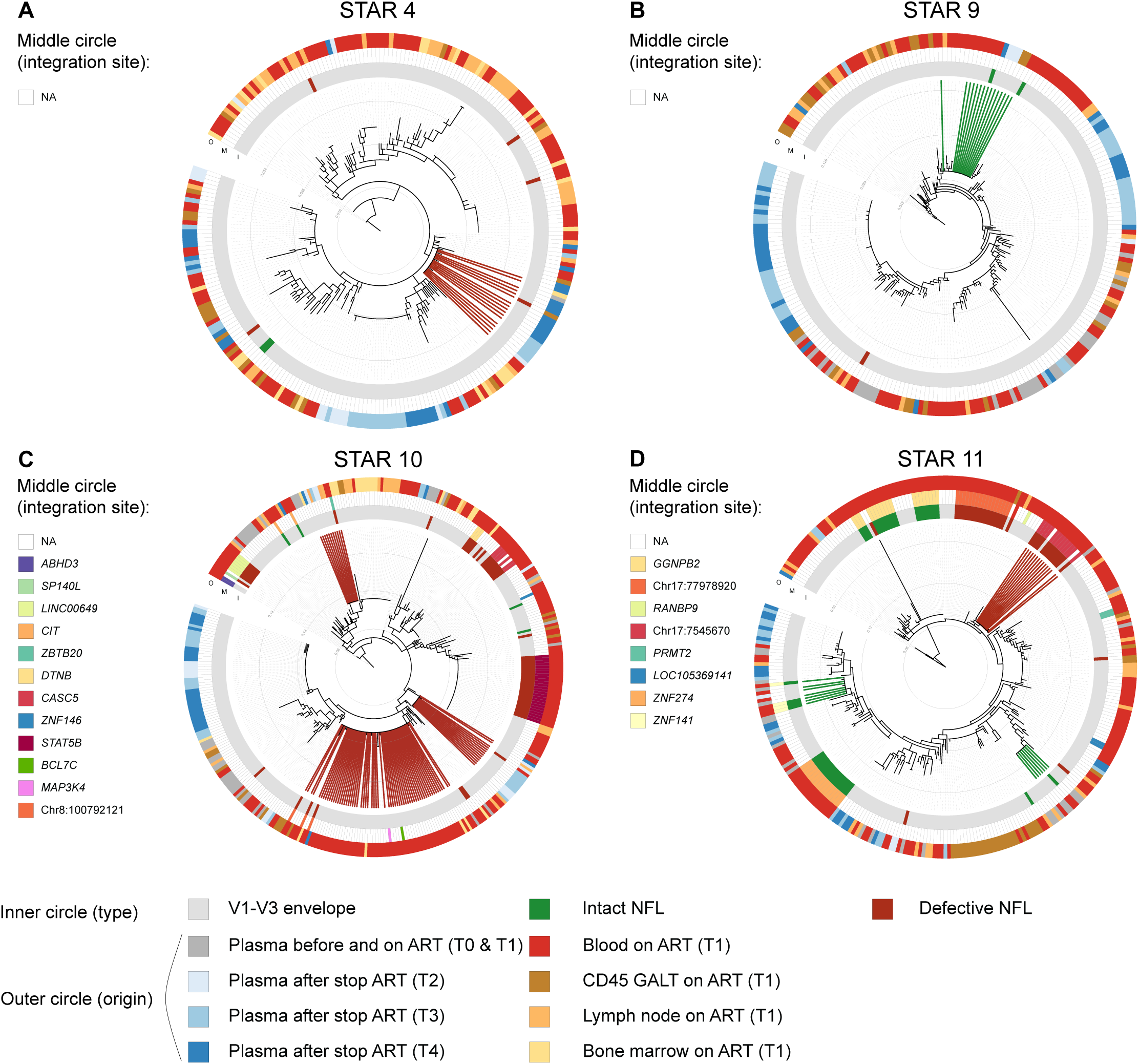
Integration sites linked to circular HIV-1 V1-V3 *env* maximum likelihood phylogenetic trees for each participant using all generated proviral and plasma V1-V3 *env* sequences before and during different stages of the ATI. The plasma and proviral sequences were obtained either prior ART initiation (timepoint 0, T0), during ART (timepoint 1, T1), 8 to 14 days after analytical treatment interruption (**ATI**) (timepoint 2, T2), at the first detectable viral load (timepoint 3, T3), and at later rebound (timepoint 4, T4). The inner circle represents the sequence type, either obtained through single-genome sequencing (**SGS**) of the V1-V3 *env* region shown in grey and V1-V3 *env* trimmed near full-length (**NFL**) genomes in colors indicating their intactness category. The middle circle shows the integration site associated with multiple displacement amplification (**MDA**) derived proviruses if available. The integration sites in the legend are shown in order of appearance on the circle. The outer circle displays the origin (sampling timepoint and/or anatomical compartment) of each plasma and proviral sequence. Matches (placed on same branch) of identical V1-V3 *env* regions between plasma and proviral NFL sequences are shown in bold lines, where the line color reflects the intactness category of the matching NFL virus. NA = not available, GALT = gut-associated lymphoid tissue.

A total of 8 matches between plasma virus and NFL proviral sequences could be found, which are highlighted corresponding to their NFL category (Figure 5, inner cicles). To further investigate the relationship between these matches in more detail, phylogenetic trees including NFL proviral sequences and V1-V3 *env* plasma sequences were constructed (Figure 6). In addition to the latter, the alignment was complemented with trimmed 3’-half viral plasma sequences obtained at T4. This analysis revealed that most of the trimmed 3’-half plasma sequences intermingled with the previously obtained V1-V3 *env* sequences, showing good accordance between the methods (Figure 6). In addition, alignments using proviral sequences and either of the 5’- or 3’- plasma viruses were constructed and checked for the potential of recombination, yet no evidence for such events was observed.

**Figure 6:**
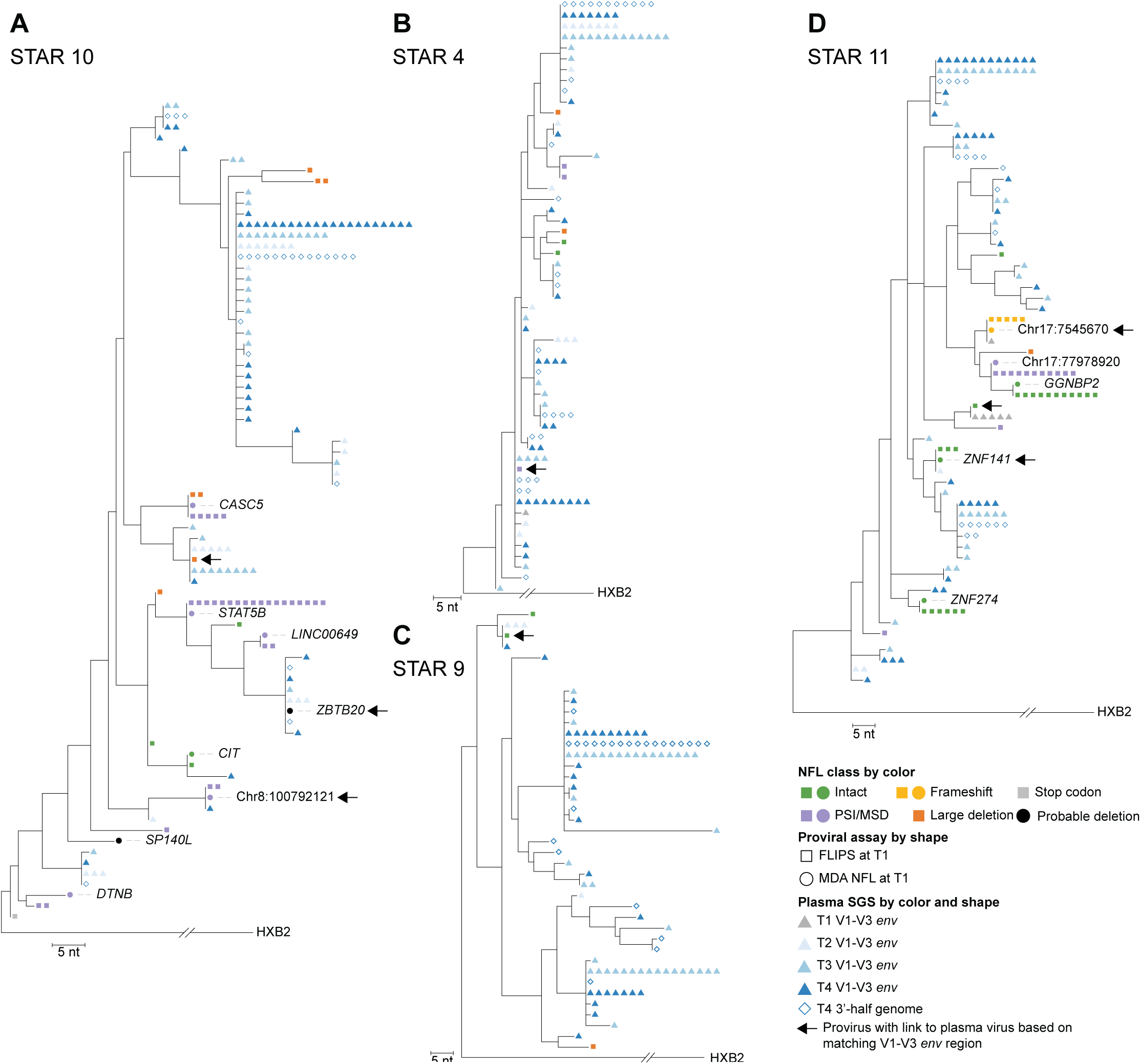
Maximum-likelihood phylogenetic trees of V1-V3 *env* sequences derived from trimmed FLIPS and/or MDA proviral sequences before ATI and (trimmed) rebounding plasma viruses before and during different stages of ATI. Proviral sequences derived from Full-Length Individual Provirus sequencing (**FLIPS**) and multiple displacement amplification (**MDA**) are shown as squares and circles respectively. The integration sites (**IS**) associated with MDA-derived proviruses are noted if available. Plasma sequences are shown as triangles (V1-V3 *env*) or diamonds (3’-half genome) where the color indicates the timepoint during analytical treatment interruption (ATI). Arrows indicate identical matches between proviral and plasma V1-V3 *env* sequences. All trees are rooted to the HXB2 reference sequence. (A) In participant STAR 10, three identical matches between defective proviral and plasma rebound sequences were found. For two, the corresponding IS *ZBTB20* and Chr8:100792121 could be recovered. (B) In participant STAR 4, only one match between a unique major splice donor (**MSD**) deleted provirus and plasma sequences was observed. (C) In STAR 9, a match between a unique intact provirus and multiple plasma sequences from different timepoints were found. (D) In STAR 11, a rebounding plasma sequence could be linked to an expansion of identical intact near full-length (**NFL**) genomes located in the *ZNF141* gene. One unique intact provirus can be linked to a residual plasma sequence from T1. SGS = single-genome sequencing,

Out of the 8 V1-V3 *env* matches between a NFL provirus and a plasma virus, 3 involved a plasma virus obtained either when ART-suppressed before the ATI (T1) or during the ATI but before a detectable viral load (T2). All of these links were found in participant STAR 11, which displays a CPS of 100% for the V1-V3 *env* region (Figure 6D). The first match was between a defective, MDA-derived provirus with a frameshift (integrated at chr17:7545670) found in both the TCM/TTM and TEM peripheral blood subsets, and a plasma virus at T1 (Figure S4D). The other matches were found between a FLIPS-derived intact provirus from the TCM/TTM peripheral blood subset and a T1 plasma virus, and an MDA-derived intact provirus from the TEM peripheral blood subset and a T2 plasma virus, respectively (Figure 6D and Figure S4D). The latter provirus was found to be integrated in the *ZNF141* gene, which belongs to the Krüppel-associated box domain (KRAB) containing zinc finger nuclease family. Interestingly, this viral sequence was not detected in the plasma from rebounding timepoints T3 and T4, however matches T0 plasma sequences (prior to ART timepoint), suggesting a phylogenetic relationship to a founder virus (Figure 5D).

Five out of the 8 plasma to proviral V1-V3 *env* matches include plasma sequences from the rebounding timepoints (T3 and T4 during ATI), but only one of them involved an intact proviral genome (Figure 6A-C). This unique FLIPS provirus in STAR 9 matched plasma sequences found at T2 prior to rebound (3/4 plasma sequences from that timepoint) and T4 during rebound. Although this provirus, located in the peripheral TEM subset, was recovered only once using FLIPS, it matched to a cluster of identical SGS V1-V3 env sequences found in cells from the peripheral blood and GALT, suggesting that it is part of a clone (Figure 5B and Figure S4B).

For participants STAR 10 and 4, one or more plasma sequences from rebound were identical to defective proviral genomes (Figure 6A-B). In fact, STAR 10 had three matches between rebounding V1-V3 *env* sequences and defective proviruses: one matched a unique FLIPS-derived provirus with a large internal deletion originating from peripheral TCM/TTM subset, one matched a provirus with a PSI/MSD deletion belonging to a clonal cell population found in the peripheral TEM subset located in an intergenic region on chromosome 8 and one matched an MDA-derived provirus from the peripheral TEM subset with a probable deletion, integrated in the *ZBTB20* gene. Note that the latter also shared a similar V1-V3 *env* sequence with two 3’-half T4 plasma sequences, yet when the alignment was extended over the entire 3’-half genome, regions with genetic differences were observed (Figure S3D). In STAR 4, a unique FLIPS-derived provirus with a PSI/MSD deletion matched to plasma V1-V3 *env* sequences from all three timepoints during the ATI (T2, T3, T4) in addition to 2 different sets of T4 3’-half genomes with identical an V1-V3 *env* (Figure 6B). This FLIPS provirus was derived from GALT, but was identical to two other V1-V3 *env* SGS sequences from the bone marrow and peripheral blood (Figure 5A). When looking at extended alignments based on either 5’- or 3’-half T4 plasma genomes and proviral sequences, a close relatedness between this provirus and the plasma half-genomes could be observed (Figure S3A). At the 5’ region, the sequences were identical beside a 105 bp deletion in the PSI/MSD region of the proviral sequence while the two sets of 3’ plasma genomes matching the V1-V3 *env* are either 8 bp or 19 bp different to the proviral sequence, suggesting a common ancestor.

In conclusion, by performing MDA-mediated NFL and IS analysis, we identified several proviruses with linked IS, predominantly belonging to peripheral blood CD4 memory subsets, that matched sequences from plasma before and/or during an ATI. Interestingly, most of these proviruses were classified as defective, raising the question whether these are still capable of producing viremia. Furthermore, some intact identical proviral sequences were detected in specimens from multiple anatomical compartments, suggesting that certain clones that harbor genome-intact proviruses can traffic between different compartments.

## Discussion

Integration of HIV-1 genomes into the DNA of host cells leads to the establishment of a persistent HIV-1 reservoir. While most of these integrated proviruses are defective, a small proportion are genetically intact and fully capable of producing infectious virions (Hiener et al., 2017). The proportion of genetically intact HIV-1 proviruses, as measured by the Intact Proviral DNA Assay (IPDA), has been shown to decay slowly, with an estimated average half-life of 4 years during the first 7 years of suppression, and 18.7 years thereafter (Peluso et al., 2020). This long half-life can in part be explained by continuous clonal expansion of infected cells harboring these genetically intact HIV-1 proviruses (Liu et al., 2020; Patro et al., 2019). While this phenomenon is well-established, the contribution of clonally expanded HIV-1 infected cells to residual viremia under ART and rebound viremia upon ATI remains underexplored. Previously, others have tried to characterize rebounding viruses by phylogenetically linking these to proviral sequences and viral sequences obtained by viral outgrowth assays (**VOA**), with limited success. While two studies were unable to find links between rebounding sequences and viral sequences recovered by VOA (Lu et al., 2018b; Vibholm et al., 2019), three other groups did find several links using similar techniques (Aamer et al., 2020; Cohen et al., 2018; Salantes et al., 2018). However, two of these latter studies were performed in the context of interventional clinical trials and the IS of these viruses remained unknown. In addition, two groups were able to link proviral sequences to rebound sequences, though only a small part of the proviral genome was queried (Barton et al., 2016; Kearney et al., 2016). We previously conducted the HIV-STAR study, where SGS on the V1-V3 *env* region was used to link proviral sequences to plasma sequences (De Scheerder et al., 2019). We found multiple links between proviral sequences and rebounding plasma sequences, however, this study was limited by the sequencing of a small subgenomic region of the proviruses and plasma viruses. In the current study, we used a combination of NFL sequencing, IS analysis and MDA-mediated IS/NFL sequencing to more accurately define the source of rebounding virus detected during ATI in a subset of HIV-STAR participants.

We first showed large discrepancies between different techniques to assess clonal expansion of HIV-1 infected cells. These discrepancies are often the result of primer biases, dictating which proviruses are amplified. This has important implications for HIV-1 reservoir research, as some assays will be unable to detect potentially relevant proviruses. In addition, we demonstrated that the use of a short subgenomic region of the HIV-1 genome (V1-V3 *env*) to assess clonality of infected cells can lead to inaccurate results. This was shown by the recovery of distinct NFL proviruses, integrated at different sites, displaying identical V1-V3 env sequences. Similar observations were made in a recently published study, where P6-PR-RT sequences were compared to matched NFL/IS sequences (Patro et al., 2019). They found multiple instances of identical proviral P6-PR-RT sequences, with distinct IS. Taken together, we conclude that evaluating clonality of HIV-1 infected cells based on the assessment of a subgenomic region should be done with caution.

We next set out to find links between NFL proviral sequences and sequences found in the plasma during different stages of an ATI. First, we identified several identical V1-V3 *env* sequences in defective proviruses and rebounding plasma viruses. Interestingly, for participant STAR 4, phylogenetic trees suggest that both V1-V3 *env* and 3’-half genome plasma SGS cluster with a provirus containing a 105 bp packaging signal deletion (including stem-loop 1 and stem-loop 2). It has been shown previously that proviruses with such defects are still capable of producing infectious virions, though with significantly lower efficiency (Pollack et al., 2017). Yet, in the 5’-half plasma dataset, a sequence was found that was completely identical to the PSI-deleted provirus in STAR 4, except that this sequence had an intact PSI, suggesting a close phylogenetic relationship.

Three other defective proviruses linked to rebound viruses, all in participant STAR 10, contain large internal deletions, making it unlikely that these are the actual source of the virus rebounding during ATI. Rather, these are probably phylogenetically related proviruses, as they share an identical V1-V3 *env* sequence. Two previous studies that tried to link proviral sequences to rebound sequences, based on full *env* sequences, concluded that while they were not able to directly link the proviral sequences to the rebounding ones, the rebounding sequences could often be accounted for by recombination (Cohen et al., 2018; Vibholm et al., 2019). Because we assessed only a small portion of the *env* gene (V1-V3 region) at timepoints T1-T3, we were not able to comprehensively study recombination events, though we hypothesize that recombination may be a probable cause of identical overlap between defective proviral sequences and rebounding virus sequences. At T4, we recovered half-genome plasma sequences, though these did not show any signs of recombination.

We further identified two links between genetically intact NFL proviruses and plasma viruses emerging upon ATI. On both occasions the intact NFL provirus was located in the peripheral TEM blood subset, suggesting these might be easier to reactivate due to higher activation status. Alternatively, this could reflect the higher degree of clonality observed in the TEM subset compared to other memory subsets, which in turn increases the chances of detecting links. The first link was found in participant STAR 9, where an intact provirus obtained with FLIPS could be linked to plasma virus at T2 and T4. Because this provirus was not retrieved in an MDA reaction, the IS remains unknown. Interestingly, this virus was first sampled at T2 and persisted into T4, which suggests that this virus emerged during the phase of an ATI when the viral load was still undetectable. In participant STAR 11, an intact provirus integrated in the *ZNF141* gene could be linked to plasma virus at T2 during an ATI. Another recent publication found a clonal infected cell population with IS in the *ZNF721/ABCA11P* gene, that contributed to persistent residual viremia which was not suppressed by ART (Halvas et al., 2020). This gene is located on chromosome 4 and belongs to the KRAB-containing zinc finger nuclease (ZNF) family. This integration event shows great similarities with the provirus we identified in the *ZNF141* gene, which also belongs to the KRAB-ZNF family and which is located on chromosome 4, just upstream of the *ZNF721/ABCA11P* gene. Interestingly, three other studies also described infected cell clones harboring a genetically-intact provirus integrated in the *ZNF721/ABCA11P* gene, suggesting that this region is a particular hotspot for the persistence of genetically intact proviruses (Einkauf et al., 2018; Halvas et al., 2020; Jiang et al., 2020). Because the plasma virus that was linked to our ZNF141 clone stems from T2, the latest timepoint with undetectable viral load during the ATI, but did not persist in the later timepoints (T3 and T4), we cannot exclude that the virus we sampled emerged as a result of continuous virus shedding, as described by Aamer *et al*. (Aamer et al., 2020), rather than ‘true’ rebounding virus. Previously, it was suggested that the origin of rebounding plasma viruses includes clonally expanded infected cells that are transcriptionally active before TI(Kearney et al., 2016). Similarly, a recent study found several overlaps between monotypic viremia that was not suppressible by ART and large proviral clones (Halvas et al., 2020). These two findings, together with the observations by Aamer *et al*. (Aamer et al., 2020), leads to the expectation that the provirus integrated in the *ZNF141* gene is a prime candidate to contribute to viral rebound, however, our current data does not support this. Off course, we cannot exclude that this viral strain was not identified at T3 and T4 because it was obscured by other rebound viruses, and thus not included among the variants we sequenced.

In a recent study it was observed that ‘elite controllers’ (EC), individuals that control HIV-1 infection spontaneously, often carry genetically intact proviral sequences integrated at spots associated with ‘deep latency’, which persist over time and are not cleared by the immune system(Jiang et al., 2020). In one EC, they described a persistently infected cell population with an intact provirus integrated in the *ZNF274* gene, which is associated with highly condensed chromatin. Interestingly, we also observed a clonally expanded infected cell population in the peripheral blood TEM fraction from STAR11, with a genetically intact provirus integrated in the *ZNF274* gene. Despite the rather large size of the clone, we did not observe the emergence of the corresponding viral sequence in the plasma during the ATI, which is in agreement with its presumed ‘deep latent’ state. In fact, it is possible that because of the heterochromatin state of the DNA at this spot, this provirus would tend to remain latent. Alternatively, we cannot exclude that this virus was not identified during the ATI due to timing of our specimen collection. Indeed, it is possible that this virus would be detected if the ATI would have been prolonged and if the participant was sampled at later time-points, especially knowing that transcription at this specific IS could be diminished and, if possible at all, would need more time to complete. These findings add to the current understanding that not all genetically intact proviruses contribute to the ‘replication competent HIV-1 viral reservoir’, as some are unlikely to rebound due to an unfavorable IS, though they may possess all the necessary attributes to rebound under specific conditions.

A study by Bertagnolli *et al*. showed that the outgrowth of a substantial fraction of viruses of the latent reservoir is blocked by autologous IgG antibodies against HIV-1 envelope (Bertagnolli et al., 2020). This mechanism might explain the discrepancy between proviruses recovered *ex vivo* and viruses recovered from the plasma. Indeed, the population of viruses that rebound might have been shaped by immune pressure, which is absent when assessing proviral sequences recovered from extracted DNA. This phenomenon further complicates finding links between proviruses and plasma viruses.

We acknowledge several limitations in this study. The first is the limited sampling from tissue compartments, possibly causing us to miss important rebound lineages. Indeed, it has been shown that tissues, including lymph nodes and GALT, harbor most of the HIV-1 latent reservoir, orders of magnitude higher than the peripheral blood compartment(Estes et al., 2017). Whether there is compartmentalization between different anatomical compartments is under debate. Several studies, including our previously conducted HIV-STAR study, have suggested that there is limited compartmentalization between the HIV-1 proviral sequences recovered from lymph nodes and from peripheral blood (Josefsson et al., 2013; Mcmanus et al., 2019; De Scheerder et al., 2019; Stockenstrom et al., 2015; Vibholm et al., 2019), based on identical proviral sequences and/or IS shared between both compartments. In addition, our previous HIV-STAR study did not show evidence of any enrichment of rebounding sequences stemming from specific anatomical compartments (De Scheerder et al., 2019), justifying our decision to focus the current study primarily on the peripheral blood compartment. The second limitation of the current study is that the link to plasma sequences at T1-T3 is based on the V1-V3 *env* region, rather than on NFL plasma sequences. This means that we cannot exclude the possibility that links between proviral sequences and rebounding plasma sequences are the result of matches in V1-V3 *env* but with genetic variation outside of this region, however the CPS for the V1-V3 *env* region for participants STAR 9 and STAR 11, which display links between intact proviral sequences and plasma rebound sequences, was calculated at 100%. Furthermore, the lack of matching sequences between half-genome plasma sequences and proviral sequences from T1 might be because at T4 the plasma typically was dominated by a genetically oligoclonal pool of viruses, which might have obscured less fit rebounding viruses that match T1 proviruses.

In conclusion, our data show that reservoir characterization using multiple methods, including IS analysis, NFL proviral sequencing and a combination of both, enables the identification of matches between proviral sequences and plasma sequences recovered before and/or during an ATI, however these matches are rare. While our findings confirm that expanded HIV infected cell clones present in the peripheral blood can contribute to both residual and rebound plasma viremia, the origins of a large fraction of rebounding viruses remained unknown. Future studies should focus on in-depth characterization of tissue reservoirs to further investigate their relative contribution to rebound viremia.

## Methods

### Samples

A total of four HIV-1 infected, ART treated participants were included in this study. All had an undetectable viral load (<20 copies/ml) for at least 1 year prior to treatment interruption, and all initiated ART during the chronic phase of infection. The participants characteristics are summarized in Table S6. Participants were sampled longitudinally, prior to and during an ATI (Figure 1B). Anatomical compartments that were sampled, and corresponding cell subsets sorted from these, are summarized in Table S1.

### CD4 T cell subset sorting

Cryopreserved PBMCs were thawed and CD4 T cell enrichment was carried out with negative magnet-activated cell sorting (Beckton Dickinson, BD IMag™, #557939). CD4 T cells were stained with the following monoclonal antibodies: CD3 (Becton Dickinson, #564465), CD8 (Becton Dickinson, #557746), CD45RO (Becton Dickinson, #555493), CD27 (Becton Dickinson, #561400), CCR7 (Becton Dickinson, #560765) and a fixable viability stain (Becton Dickinson, #565388). Fluorescence-activated cell sorting was used to sort stained peripheral blood-derived CD4 T cells into naïve CD4 T cells (CD45RO-, CD45RA+), central memory/transitional memory CD4 T cells (CD3+ CD8-CD45RO+ CD27) and effector memory CD4 T cells (CD3+ CD8-CD45RO+ CD27-), GALT cells into CD45+ cells and cells from lymph nodes into central memory/transitional memory CD4 T cells (CD3+ CD8-CD45RO+ CD27+) and effector memory CD4 T cells (CD3+ CD8-CD45RO+ CD27-), using a BD FACSJazz cell sorter machine, as previously described (De Scheerder et al., 2019). The gating strategy used for the aforementioned sorts can be found in Figure S5. A small fraction of each sorted cell population was analyzed by flow cytometry to check for purity, which was over 95% on average. Flow cytometry data was analyzed using FlowJo software (Tree-Star).

### Droplet digital PCR (ddPCR)

Sorted cells were pelleted and lysed in 100µL lysis buffer (10mM TrisHCl, 0.5% NP-40, 0.5% Tween-20 and proteinase K at 20 mg/ml) by incubating for 1 hour at 55°C and 15 min at 85°C. HIV-1 copy number was determined by a total HIV-1 DNA assay on droplet digital PCR (Bio-Rad, QX200 system), as described previously (Rutsaert et al., 2019). PCR amplification was carried out with the following cycling program: 10 min at 98°C; 45 cycles (30 sec at 95°C, 1 min at 58°C); 10 min at 98°C. Droplets were read on a QX200 droplet reader (Bio-Rad). Analysis was performed using ddpcRquant software (Trypsteen et al., 2015).

### Whole genome amplification (WGA)

Cell lysates were diluted according to ddPCR HIV-1 copy quantification, so that less than 30% of reactions contained a single proviral genome. Whole genome amplification was performed by multiple displacement amplification with the REPLI-g single cell kit (Qiagen, #150345), according to manufacturer’s instructions. The resulting amplification product was split for downstream IS analysis, single genome/proviral sequencing, and, for selected reactions, near full-length HIV-1 sequencing.

### Single genome/proviral sequencing

Single genome/proviral sequencing (SGS) of the V1-V3 region of env was performed as described before (Josefsson et al., 2012; Von Stockenstrom et al., 2015), with a few adaptations. The amplification consists of a nested PCR with the following primers: Round 1 forward (E20) 5’-GGGCCACACATGCCTGTGTACCCACAG-3’ and reverse (E115) 5’-AGAAAAATTCCCCTCCACAATTAA-3’; round 2, forward (E30) 5’-GTGTACCCACAGACCCCAGCCCACAAG-3’ and reverse (E125) 5’-CAATTTCTGGGTCCCCTCCTGAGG-3’. The 25 µL PCR mix for the first round is composed of: 5 µL 5X Mytaq buffer, 0.375 µL Mytaq polymerase (Bioline, #BIO-21105), 400 nM forward primer, 400 nM reverse primer and 1 µL REPLI-g product. The mix for the second round has the same composition and takes 1 µL of the first-round product as an input. Thermocycling conditions for first and second PCR rounds are as follows: 2 min at 94°C; 35 cycles (30 sec at 94°C, 30 sec at 60°C, 1 min at 72°C); 5 min at 72°C. Resulting amplicons were visualized on a 1% agarose gel and Sanger sequenced (Eurofins Genomics, Ebersberg, Germany) from both ends, using second round PCR primers. Both 5’- and 3’-half genome amplicons were generated from T4 plasma samples. RNA was extracted from the virions and cDNA was generated as follows: 1) Plasma samples were thawed at 37°C. 2) Remove debris by centrifuging the plasma for 10 min at 3600 rpm and discarding the pellet. 3) Transfer supernatant to ultracentrifuge tube and adjust volume to 9 mL with PBS. 4) Centrifuge at

>85.000 g for 70 min at 4°C. 5) 240 µL of the supernatant is subjected to viral RNA extraction using the QIAamp Viral RNA Mini Kit (Qiagen, #52904), according to manufacturer’s instructions. 6) Half of the RNA was used to generated cDNA for 5’-half sequencing using the R5968 primer (5’-TGTCTYCKCTTCTTCCTGCCATAG-3’), while the other half was used to generate cDNA for 3’-half sequencing using the primer R9665 (5’-GTCTGAGGGATCTCTAGWTACCAGA-3’). Two mastermixes were prepared. Mix 1 consisted of 25 µL RNA, 2.5 µL 10mM dNTP, 1.25 µL 20 µM oligo-dT (SuperScript III First Strand synthesis system, Invitrogen, #18080051) and 0.5 µL 50 µM primer. Mix 2 consisted of 0.75 µL RNAse free water, 5X RT buffer (Invitrogen, #18080051), 2.5 µL 100 mM DTT, 2.5 µL 40U/ µL RNAse inhibitor (Takara, #2313B), 2.5 µL SuperScript III Reverse Transcriptase (Invitrogen, #18080051) and 2 µL ThermaSTOP RT (Sigma Aldrich, #TSTOPRT-250). Mix 1 was heated to 65°C for 5 min and then snap-chilled on ice for at least 2 min. Mix 2 was pre-warmed to 50°C and then added to the chilled mix 1. The mixture was incubated at 50°C for 90 min. 1 µL SuperScript III Reverse Transcriptase was added to the reaction, followed by another incubation of 90 min at 50°C and then 70°C for 15 min. Finally, 1 µL RNAse H (Invitrogen, #18080051) was added, followed by an incubation of 20 min at 37°C. Subsequently, the cDNA was used as template for half-genome long-range PCRs, as described previously (Cole et al., 2021). The 25 µL PCR mix for the first round was composed of: 5 µL 5X Prime STAR GXL buffer, 0.5 µL PrimeStar GXL polymerase (Takara Bio, #R050B), 0.125 µL ThermaStop (Sigma Aldrich, #TSTOP-500), 250 nM forward and reverse primers, and 1 µL MDA product. The mix for the second round had the same composition and took 1 µL of the first-round product as an input. Thermocycling conditions for first and second PCR rounds were as follows: 2 min at 98°C; 35 cycles (10 sec at 98°C, 15 sec at 62°C, 5 min at 68°C); 7 min at 68°C. Reactions without reverse transcriptase were negative, ensuring that the RNA extracts were not contaminated by DNA. PCR products were checked on a 1% agarose gel and positives were sequenced by Illumina sequencing, as described below.

### Integration site loop amplification (ISLA)

Integration site sequencing was carried out by integration site loop amplification (ISLA), as described by Wagner *et al*. (Wagner et al., 2014a), but with a few modifications. Firstly, the *env* primer used during the linear amplification step was omitted, as it was not necessary to recover the *env* portion of the provirus at a later stage. Therefore, the reaction was not split after the linear amplification, and the entire reaction was used as an input into subsequent decamer binding and loop formation. For some proviruses, an alternative set of primers were used to retrieve the IS from the 5’ end (Table S7). Resulting amplicons were visualized on a 1% agarose gel and positives were sequenced by Sanger sequencing. Analysis of the generated sequences was performed using the ‘Integration Sites’ webtool developed by the Mullins lab; https://indra.mullins.microbiol.washington.edu/integrationsites/.

### Full-length individual proviral sequencing assay

Proviral sequences from the genomic DNA of sorted subsets were recovered by the Full-length Individual Proviral Sequencing (FLIPS) assay as first described by Hiener *et al*. (Hiener et al., 2017) with some minor alterations. Briefly, the assay consists of two rounds of nested PCR at an end-point dilution where 30% of the wells are positive. This yields proviral fragments of up to 9 kb using the following primers for the first round BLOuterF (5’-AAATCTCTAGCAGTGGCGCCCGAACAG-3’) and BLOuterR (5’-TGAGGGATCTCTAGTTACCAGAGTC-3’) followed by a second round using primers 275F (5’-ACAGGGACCTGAAAGCGAAAG-3’) and 280R (5’-CTAGTTACCAGAGTCACACAACAGACG-3’). The cycling conditions are 94°C for 2 m; then 94°C for 30 s, 64°C for 30 s, 68°C for 10 m for 3 cycles; 94°C for 30 s, 61°C for 30 s, 68°C for 10 m for 3 cycle; 94°C for 30 s, 58°C for 30 s, 68°C for 10 m for 3 cycle; 94°C for 30 s, 55°C for 30 s, 68°C for 10 m for 21 cycle; then 68°C for 10 m. For the second round, 10 extra cycles at 55°C are included. The PCR products were visualized using agarose gel electrophoresis. Amplified proviruses from positive wells were cleaned using AMPure XP beads (Beckman Coulter), followed by a quantification of each cleaned provirus with Quant-iT PicoGreen dsDNA Assay Kit (Invitrogen). Next, an NGS library preparation using the Nextera XT DNA Library Preparation Kit (Illumina) with indexing of 96-samples per run was used according to the manufacturer’s instructions, except that input and reagents volumes were halved and libraries were normalized manually. The pooled library was sequenced on a MiSeq Illumina platform via 2×150 nt paired-end sequencing using the 300 cycle v2 kit.

### Near full-length provirus amplification from MDA reactions

MDA reactions containing a potentially clonal proviral sequence were subjected to near full-length proviral sequencing, using either a single-amplicon approach (Hiener et al., 2017), a four-amplicon approach (Patro et al., 2019), or a five-amplicon approach (Einkauf et al., 2018), as previously described. In case of the multiple amplicon approaches, amplicons were pooled equimolarly and sequenced as described above.

### *De Novo* assembly of HIV-1 proviruses and analysis

The generated sequencing data from either FLIPS or multiple amplicon approaches was demultiplexed and used to *de novo* assemble individual proviruses. The code used to perform *de novo* assembly can be found at the following GitHub page: https://github.com/laulambr/virus_assembly. In short, the workflow consists of following steps: (i) check of sequencing quality for each library using FastQC (http://www.bioinformatics.babraham.ac.uk/projects/fastqc) and removal of Illumina adaptor sequences and trimming of 5’ and 3’ terminal ends using BBtools (sourceforge.net/projects/bbmap/). (ii) The trimmed reads are *de novo* assembled using MEGAHIT (Li et al., 2015) generating contigs for each library. (iii) Per library, all *de novo* contigs were checked using blastn against the HXB2 reference virus as a filter to exclude non-HIV-1 contigs in the following analysis steps. (iv) Subsequently, the trimmed reads were mapped against the *de novo* assembled HIV-1 contigs to enable the calling of the final majority consensus sequence of each provirus using bbmap. Alignments of proviral sequences for each participant were made via MAFFT (Katoh et al., 2002) and manually inspected via MEGA7 (Kumar et al., 2016). The generated HIV-1 proviruses were categorized as intact or defective as described previously (Hiener et al., 2017). NFL proviruses and half-genome plasma sequences were screened for recombination by the “DualBrothers” software (Minin et al., 2005) and the “Recombinant Identification Program” webtool from the Los Alamos National Laboratory HIV sequence database (https://www.hiv.lanl.gov). Phylogenetic trees were constructed using PhyML v3.0 (Guindon et al., 2010) (best of NNI and SPR rearrangements) and 1000 bootstraps. MEGA7 (Kumar et al., 2016) and iTOL v5 (Letunic and Bork, 2019) were used to visualize phylogenetic trees.

### Statistical analysis

P-values in Figure 2A and Figure 2C test for a difference in the proportion of respectively unique IS or unique proviruses between TCM/TTM and TEM subsets. P-values were calculated using “prop.test” command in R versions 3.6.2 (“R Core Team,” 2020). Infection frequencies for FLIPS data were calculated by expressing the total number of identified HIV positive cells as a proportion of all cells analysed. The infection frequency was compared across cellular subsets using a logistic regression on the number of cells positive for HIV and total number of cells using “glm” function in R. Interaction between participant and cellular subset was detected (P < 0.001) and included in the logistic regression. P-values were calculated using the “Anova” function from the “car” package in R(John and Sanford, 2019).

## Data availability statement

Data will be uploaded to public repositories upon acceptance of the manuscript.

## Study approval

This study was approved by the Ethics Committee of the Ghent University Hospital (Belgian registration number: B670201525474). Written informed consent was obtained from all study participants.

## Acknowledgements and funding sources

We would like to acknowledge and thank all participants who donated samples to the HIV-STAR study, and all the MDs and study nurses that assisted with the sample collection. We would also like to thank Marion Pardons, Tine Struyve and Sofie Rutsaert for providing guidance during initial data analyses, for the constructive discussions and for critically reading the manuscript. We are grateful for the discussions with and input from James Mullins, Rafick Sékaly, Susan Pereira Ribeiro, Hadega Aamer, Sam Kint, Oleg Denisenko, Katie Fisher and Bethany Horsburgh. In addition, we would like to thank Kim De Leeneer, Céline Helsmoortel and Bram Parton for their assistance in performing MiSeq sequencing at UZ Ghent. This current research work was supported by the NIH (R01-AI134419, MPI: LV and LF) and the Research Foundation Flanders (S000319N and G0B3820N). LV was supported by the Research Foundation Flanders (1.8.020.09.N.00) and the Collen-Francqui Research Professor Mandate. SP was supported by the Delaney AIDS Research Enterprise (DARE) to Find a Cure (1U19AI096109 and 1UM1AI126611-01) and the Australian National Health and Medical Research Council (APP1061681 and APP1149990). The sample collection at UZ Ghent was supported by an MSD investigator grant (ISS 52777). BC and LL were supported by FWO Vlaanderen (1S28918N, 1S29220N). BV was supported by a postdoctoral grant (12U7121N) of the Research Foundation - Flanders (Fonds voor Wetenschappelijk Onderzoek).

## Author contributions

BC, LL, LF, SP and LV conceptualized the experiments. MADS processed the samples from the initial HIV STAR study, including cell isolation from peripheral blood and tissue, and she performed cell sorting and single-genome sequencing. BC and YN performed experiments involving cell sorting, multiple displacement amplification, single-genome sequencing and integration site sequencing. LL and ZB performed experiments involving near full-length proviral sequencing. BC, LL, BV, JSE and TS analyzed data and performed associated analyses. BC, LL, TS and BV made figures and tables. BC and LL wrote the manuscript. All co-authors edited and approved the manuscript.

## References

Aamer, H.A., Mcclure, J., Ko, D., Maenza, J., Collier, A.C., Mullins, J.I., and Frenkel, L.M. (2020). Cells producing residual viremia during antiretroviral treatment appear to contribute to rebound viremia following interruption of treatment. PLOS Pathogens 16, e1008791.

Artesi, M., Hahaut, V., Cole, B., Lambrechts, L., Ashrafi, F., Marçais, A., Hermine, O., Griebel, P., Arsic, N., van der Meer, F., et al. (2021). PCIP-seq: simultaneous sequencing of integrated viral genomes and their insertion sites with long reads. Genome Biol 22, 97.

Bailey, J.R., Sedaghat, A.R., Kieffer, T., Brennan, T., Lee, P.K., Wind-rotolo, M., Haggerty, C.M., Kamireddi, A.R., Liu, Y., Lee, J., et al. (2006). Residual Human Immunodeficiency Virus Type 1 Viremia in Some Patients on Antiretroviral Therapy Is Dominated by a Small Number of Invariant Clones Rarely Found in Circulating CD4+ T Cells. Journal of Virology 80, 6441–6457.

Barton, K., Hiener, B., Winckelmann, A., Rasmussen, T.A., Shao, W., Byth, K., Lanfear, R., Solomon, A., Mcmahon, J., Harrington, S., et al. (2016). Broad activation of latent HIV-1 in vivo. Nature Communications 7, 1–8.

Bertagnolli, L.N., Varriale, J., Sweet, S., Brockhurst, J., Simonetti, F.R., White, J., Beg, S., Lynn, K., Mounzer, K., Frank, I., et al. (2020). Autologous IgG antibodies block outgrowth of a substantial but variable fraction of viruses in the latent reservoir for HIV-1. Proc Natl Acad Sci U S A 117, 32066–32077.

Boritz, E.A., Darko, S., Swaszek, L., Wolf, G., Wells, D., Wu, X., Henry, A.R., Laboune, F., Hu, J., Ambrozak, D., et al. (2016). Multiple Origins of Virus Persistence during Natural Control of HIV Infection. Cell 166, 1004–1015.

Brennan, T.P., Woods, J.O., Sedaghat, A.R., Siliciano, J.D., Siliciano, R.F., and Wilke, C.O. (2009). Analysis of Human Immunodeficiency Virus Type 1 Viremia and Provirus in Resting CD4+ T Cells Reveals a Novel Source of Residual Viremia in Patients on Antiretroviral Therapy. Journal of Virology 83, 8470–8481.

Cesana, D., Santoni de Sio, F.R., Rudilosso, L., Gallina, P., Calabria, A., Beretta, S., Merelli, I., Bruzzesi, E., Passerini, L., Nozza, S., et al. (2017). HIV-1-mediated insertional activation of STAT5B and BACH2 trigger viral reservoir in T regulatory cells. Nature Communications 8, 498.

Chun, T., and Fauci, A.S. (1999). Perspective Latent reservoirs of HIV[]: Obstacles to the eradication of virus. Proceedings of the National Academy of Sciences of the United States of America 96, 10958–10961.

Chun, T.W., Stuyver, L., Mizell, S.B., Ehler, L. a, Mican, J. a, Baseler, M., Lloyd, a L., Nowak, M. a, and Fauci, a S. (1997). Presence of an inducible HIV-1 latent reservoir during highly active antiretroviral therapy. Proceedings of the National Academy of Sciences of the United States of America 94, 13193–13197.

Chun, T.W., Engel, D., Berrey, M.M., Shea, T., Corey, L., and Fauci, A.S. (1998). Early establishment of a pool of latently infected, resting CD4(+) T cells during primary HIV-1 infection. Proceedings of the National Academy of Sciences of the United States of America 95, 8869–8873.

Clarridge, K.E., Blazkova, J., Einkauf, K., Petrone, M., Refsland, W., Justement, J.S., Shi, V., Huiting, E.D., Seamon, C.A., Lee, G.Q., et al. (2018). Effect of analytical treatment interruption and reinitiation of antiretroviral therapy on HIV reservoirs and immunologic parameters in infected individuals. PLoS Pathogens 14, e1006792.

Cohen, Y.Z., Lorenzi, J.C.C., Krassnig, L., Barton, J.P., Burke, L., Pai, J., Lu, C.L., Mendoza, P., Oliveira, T.Y., Sleckman, C., et al. (2018). Relationship between latent and rebound viruses in a clinical trial of anti – HIV-1 antibody 3BNC117. Journal of Experimental Medicine 215, 2311–2324.

Cohn, L.B., Silva, I.T., Oliveira, T.Y., Rosales, R.A., Parrish, E.H., Learn, G.H., Hahn, B.H., Czartoski, J.L., McElrath, M.J., Lehmann, C., et al. (2015a). HIV-1 integration landscape during latent and active infection. Cell 160, 420–432.

Cohn, L.B., Silva, I.T., Oliveira, T.Y., Rosales, R.A., Parrish, E.H., Learn, G.H., Hahn, B.H., Czartoski, J.L., McElrath, M.J., Lehmann, C., et al. (2015b). HIV-1 integration landscape during latent and active infection. Cell 160, 420–432.

Cole, B., Lambrechts, L., Gantner, P., Noppe, Y., Bonine, N., Witkowski, W., Chen, L., Palmer, S., Mullins, J.I., Chomont, N., et al. (2021). In-depth single-cell analysis of translation-competent HIV-1 reservoirs identifies cellular sources of plasma viremia. Nat Commun 12, 3727.

Einkauf, K., Lee, G.Q., Gao, C., Sharaf, R., Sun, X., Hua, S., Chen, S., Jiang, C., Lian, X., Chowdhury, F.Z., et al. (2018). Distinct chromosomal positions of intact HIV-1 proviruses. Journal of Clinical Investigation 129, 988–998.

Estes, J.D., Kityo, C., Ssali, F., Swainson, L., Makamdop, K.N., Prete, G.Q. Del, Deeks, S.G., Luciw, P.A., Chipman, J.G., Beilman, G.J., et al. (2017). Defining total-body AIDS-virus burden with implications for curative strategies. Nature Medicine 23, 1271–1276.

Finzi, D., Hermankova, M., Pierson, T., Carruth, L.M., Buck, C., Chaisson, R.E., Quinn, T.C., Chadwick, K., Margolick, J., Brookmeyer, R., et al. (1997). Identification of a reservoir for HIV-1 in patients on highly active antiretroviral therapy. Science 278, 1295–1300.

Garner, S.A., Rennie, S., Ananworanich, J., Dube, K., Margolis, D.M., Sugarman, J., Tressler, R., Gilbertson, A., and Dawson, L. (2017). Interrupting antiretroviral treatment in HIV cure research[]: scientific and ethical considerations. Journal of Virus Eradication 3, 82–84.

Guindon, S., Dufayard, J.-F., Lefort, V., Anisimova, M., Hordijk, W., and Gascuel, O. (2010). New Algorithms and Methods to Estimate Maximum-Likelihood Phylogenies: Assessing the Performance of PhyML 3.0. Systematic Biology 59, 307–321.

Halvas, E.K., Hughes, S.H., Mellors, J.W., Halvas, E.K., Joseph, K.W., Brandt, L.D., Guo, S., Sobolewski, M.D., Jacobs, J.L., Tumiotto, C., et al. (2020). HIV-1 viremia not suppressible by antiretroviral therapy can originate from large T cell clones producing infectious virus. Journal of Clinical Investigation 130, 5847–5857.

Hiener, B., Horsburgh, B.A., Eden, J.S., Barton, K., Schlub, T.E., Lee, E., von Stockenstrom, S., Odevall, L., Milush, J.M., Liegler, T., et al. (2017). Identification of Genetically Intact HIV-1 Proviruses in Specific CD4+T Cells from Effectively Treated Participants. Cell Reports 21, 813–822.

Hosmane, N.N., Kwon, K.J., Bruner, K.M., Capoferri, A.A., Beg, S., Rosenbloom, D.I.S., Keele, B.F., Ho, Y.-C., Siliciano, J.D., and Siliciano, R.F. (2017). Proliferation of latently infected CD4 + T cells carrying replication-competent HIV-1: Potential role in latent reservoir dynamics. The Journal of Experimental Medicine 214, 959–972.

Jiang, C., Lian, X., Gao, C., Sun, X., Einkauf, K.B., Chevalier, J.M., Chen, S.M.Y., Hua, S., Rhee, B., Chang, K., et al. (2020). Distinct viral reservoirs in individuals with spontaneous control of HIV-1. Nature 585, 261–267.

John, F., and Sanford, W. (2019). An R Companion to Applied Regression (Thousand Oaks CA: Sage).

Josefsson, L., Eriksson, S., Sinclair, E., Ho, T., Killian, M., Epling, L., Shao, W., Lewis, B., Bacchetti, P., Loeb, L., et al. (2012). Hematopoietic Precursor Cells Isolated From Patients on Long-term Suppressive HIV Therapy Did Not Contain HIV-1 DNA. Journal of Infectious Diseases 206, 28–34.

Josefsson, L., von Stockenstrom, S., Faria, N.R., Sinclair, E., Bacchetti, P., Killian, M., Epling, L., Tan, A., Ho, T., Lemey, P., et al. (2013). The HIV-1 reservoir in eight patients on long-term suppressive antiretroviral therapy is stable with few genetic changes over time. Proceedings of the National Academy of Sciences of the United States of America 110, E4987–96.

Katoh, K., Misawa, K., Kuma, K., and Miyata, T. (2002). MAFFT: a novel method for rapid multiple sequence alignment based on fast Fourier transform. Nucleic Acids Research 30, 3059–3066.

Kearney, M.F., Wiegand, A., Shao, W., Coffin, J.M., Mellors, J.W., Lederman, M., Gandhi, R.T., Keele, B.F., and Li, J.Z. (2016). Origin of Rebound Plasma HIV Includes Cells with Identical Proviruses That Are Transcriptionally Active before Stopping of Antiretroviral Therapy. Journal of Virology 90, 1369–1376.

Kumar, S., Stecher, G., and Tamura, K. (2016). MEGA7: Molecular Evolutionary Genetics Analysis Version 7.0 for Bigger Datasets. Molecular Biology and Evolution 33, 1870–1874.

Lambrechts, L., Cole, B., Rutsaert, S., Trypsteen, W., and Vandekerckhove, L. (2020). Emerging PCR-Based Techniques to Study HIV-1 Reservoir Persistence. Viruses 12, 1–12.

Laskey, S.B., Pohlmeyer, C.W., Bruner, K.M., and Siliciano, R.F. (2016). Evaluating Clonal Expansion of HIV-Infected Cells: Optimization of PCR Strategies to Predict Clonality. PLOS Pathogens 12, e1005689.

Lee, G.Q., Orlova-Fink, N., Einkauf, K., Chowdhury, F.Z., Sun, X., Harrington, S., Kuo, H.-H., Hua, S., Chen, H.-R., Ouyang, Z., et al. (2017). Clonal expansion of genome-intact HIV-1 in functionally polarized Th1 CD4+ T cells. Journal of Clinical Investigation 127, 2689–2696.

Letunic, I., and Bork, P. (2019). Interactive Tree Of Life (iTOL) v4: recent updates and new developments. Nucleic Acids Research 47, W256–W259.

Li, D., Liu, C.-M., Luo, R., Sadakane, K., and Lam, T.-W. (2015). MEGAHIT: an ultra-fast single-node solution for large and complex metagenomics assembly via succinct de Bruijn graph. Bioinformatics 31, 1674–1676.

Liu, R., Simonetti, F.R., and Ho, Y. (2020). The forces driving clonal expansion of the HIV-1 latent reservoir. Virology Journal 17.

Lu, C., Pai, J.A., Nogueira, L., Mendoza, P., Gruell, H., Oliveira, T.Y., and Barton, J. (2018a). Relationship between intact HIV-1 proviruses in circulating CD4 + T cells and rebound viruses emerging during treatment interruption. Proceedings of the National Academy of Sciences of the United States of America 115, 11341–11348.

Lu, C.-L., Pai, J.A., Nogueira, L., Mendoza, P., Gruell, H., Oliveira, T.Y., Barton, J., Lorenzi, J.C.C., Cohen, Y.Z., Cohn, L.B., et al. (2018b). Relationship between intact HIV-1 proviruses in circulating CD4 + T cells and rebound viruses emerging during treatment interruption. Proceedings of the National Academy of Sciences of the United States of America 115, 11341–11348.

Maldarelli, F., Wu, X., Su, L., Simonetti, F.R., Shao, W., Hill, S., Spindler, J., Ferris, A.L., Mellors, J.W., Kearney, M.F., et al. (2014). Specific HIV integration sites are linked to clonal expansion and persistence of infected cells. Science 345, 179–183.

Mcmanus, W.R., Coffin, J.M., Kearney, M.F., Mcmanus, W.R., Bale, M.J., Spindler, J., Wiegand, A., Musick, A., Patro, S.C., Sobolewski, M.D., et al. (2019). HIV-1 in lymph nodes is maintained by cellular proliferation during antiretroviral therapy. Journal of Clinical Investigation 129, 4629–4642.

Minin, V.N., Dorman, K.S., Fang, F., and Suchard, M.A. (2005). Dual multiple change-point model leads to more accurate recombination detection. Bioinformatics 21, 3034–3042.

Pannus, P., Rutsaert, S., De Wit, S., Allard, S.D., Vanham, G., Cole, B., Nescoi, C., Aerts, J., De Spiegelaere, W., Tsoumanis, A., et al. (2020). Rapid viral rebound after analytical treatment interruption in patients with very small HIV reservoir and minimal on-going viral transcription. Journal of the International AIDS Society 23, e25453.

Patro, S.C., Brandt, L.D., Bale, M.J., Halvas, E.K., Joseph, K.W., Shao, W., and Wu, X. (2019). Combined HIV-1 sequence and integration site analysis informs viral dynamics and allows reconstruction of replicating viral ancestors. Proceedings of the National Academy of Sciences of the United States of America 116, 25891–25899.

Peluso, M.J., Laird, G.M., Deeks, S.G., Peluso, M.J., Bacchetti, P., Ritter, K.D., Beg, S., Lai, J., Martin, J.N., Hunt, P.W., et al. (2020). Differential decay of intact and defective proviral DNA in HIV-1 – infected individuals on suppressive antiretroviral therapy. JCI Insight 5, e132997.

Pinzone, M.R., Vanbelzen, D.J., Weissman, S., Bertuccio, M.P., Cannon, L., Venanzi-rullo, E., Migueles, S., Jones, R.B., Mota, T., Joseph, S.B., et al. (2019). Longitudinal HIV sequencing reveals reservoir expression leading to decay which is obscured by clonal expansion. Nature Communications 10.

Pollack, R.A., Jones, R.B., Pertea, M., Bruner, K.M., Martin, A.R., Thomas, A.S., Capoferri, A.A., Beg, S.A., Huang, S.H., Karandish, S., et al. (2017). Defective HIV-1 Proviruses Are Expressed and Can Be Recognized by Cytotoxic T Lymphocytes, which Shape the Proviral Landscape. Cell Host and Microbe 21, 494–506.

“R Core Team” (2020). R: A language and environment for statistical computing.

Rutsaert, S., Spiegelaere, W. De, Clercq, L. De, and Vandekerckhove, L. (2019). Evaluation of HIV-1 reservoir levels as possible markers for virological failure during boosted darunavir monotherapy. Journal of Antimicrobial Chemotherapy 74, 3030–3034.

Salantes, D.B., Tebas, P., Bar, K.J., Salantes, D.B., Zheng, Y., Mampe, F., Srivastava, T., Beg, S., Lai, J., Li, J.Z., et al. (2018). HIV-1 latent reservoir size and diversity are stable following brief treatment interruption. Journal of Clinical Investigation 128, 3102–3115.

De Scheerder, M.-A., Vrancken, B., Dellicour, S., Schlub, T., Lee, E., Shao, W., Rutsaert, S., Verhofstede, C., Kerre, T., Malfait, T., et al. (2019). HIV Rebound Is Predominantly Fueled by Genetically Identical Viral Expansions from Diverse Reservoirs. Cell Host & Microbe 26, 347–358.

Simonetti, F.R., Sobolewski, M.D., Fyne, E., Shao, W., Spindler, J., Hattori, J., Anderson, E.M., Watters, S.A., Hill, S., Wu, X., et al. (2016). Clonally expanded CD4+ T cells can produce infectious HIV-1 in vivo. Proceedings of the National Academy of Sciences of the United States of America 113, 1883–1888.

Stockenstrom, S. Von, Odevall, L., Lee, E., Sinclair, E., Bacchetti, P., Killian, M., Epling, L., Shao, W., Hoh, R., Ho, T., et al. (2015). Longitudinal Genetic Characterization Reveals That Cell Proliferation Maintains a Persistent HIV Type 1 DNA Pool During Effective HIV Therapy. Journal of Infectious Diseases 212, 596–607.

Von Stockenstrom, S., Odevall, L., Lee, E., Sinclair, E., Bacchetti, P., Killian, M., Epling, L., Shao, W., Hoh, R., Ho, T., et al. (2015). Longitudinal Genetic Characterization Reveals That Cell Proliferation Maintains a Persistent HIV Type 1 DNA Pool during Effective HIV Therapy. Journal of Infectious Diseases 212, 596–607.

Tobin, N.H., Learn, G.H., Holte, S.E., Wang, Y., Melvin, A.J., McKernan, J.L., Pawluk, D.M., Mohan, K.M., Lewis, P.F., Mullins, J.I., et al. (2005). Evidence that low-level viremias during effective highly active antiretroviral therapy result from two processes: expression of archival virus and replication of virus. Journal of Virology 79, 9625–9634.

Trypsteen, W., Vynck, M., Neve, J. De, Bonczkowski, P., Vandekerckhove, L., Spiegelaere, W. De, De Neve, J., Bonczkowski, P., Kiselinova, M., Malatinkova, E., et al. (2015). ddpcRquant: threshold determination for single channel droplet digital PCR experiments. Analytical and Bioanalytical Chemistry 407, 5827–5834.

Vibholm, L.K., Lorenzi, J.C.C., Pai, J.A., Cohen, Y.Z., Oliveira, T.Y., Barton, J.P., Garcia Noceda, M., Lu, C.-L., Ablanedo-Terrazas, Y., Del Rio Estrada, P.M., et al. (2019). Characterization of Intact Proviruses in Blood and Lymph Node from HIV-Infected Individuals Undergoing Analytical Treatment Interruption. Journal of Virology 93, e01920–18.

Wagner, T.A., McKernan, J.L., Tobin, N.H., Tapia, K.A., Mullins, J.I., Frenkel, M., and Frenkel, L.M. (2013). An increasing proportion of monotypic HIV-1 DNA sequences during antiretroviral treatment suggests proliferation of HIV-infected cells. Journal of Virology 87, 1770–1778.

Wagner, T.A., McLaughlin, S., Garg, K., Cheung, C.Y.K., Larsen, B.B., Styrchak, S., Huang, H.C., Edlefsen, P.T., Mullins, J.I., and Frenkel, L.M. (2014a). Proliferation of cells with HIV integrated into cancer genes contributes to persistent infection. Science 345, 570–573.

Wagner, T.A., McLaughlin, S., Garg, K., Cheung, C.Y.K., Larsen, B.B., Styrchak, S., Huang, H.C., Edlefsen, P.T., Mullins, J.I., and Frenkel, L.M. (2014b). Proliferation of cells with HIV integrated into cancer genes contributes to persistent infection. Science 345, 570–573.

Wang, Z., Gurule, E.E., Brennan, T.P., Gerold, J.M., Kwon, K.J., Hosmane, N.N., Kumar, M.R., Beg, S.A., Capoferri, A.A., Ray, S.C., et al. (2018). Expanded cellular clones carrying replication-competent HIV-1 persist, wax, and wane. Proceedings of the National Academy of Sciences of the United States of America 115, E2575–E2584.

