## Supplementary Information for "Extensive characterization of HIV-1 reservoirs reveals links to plasma viremia before and during analytical treatment interruption"

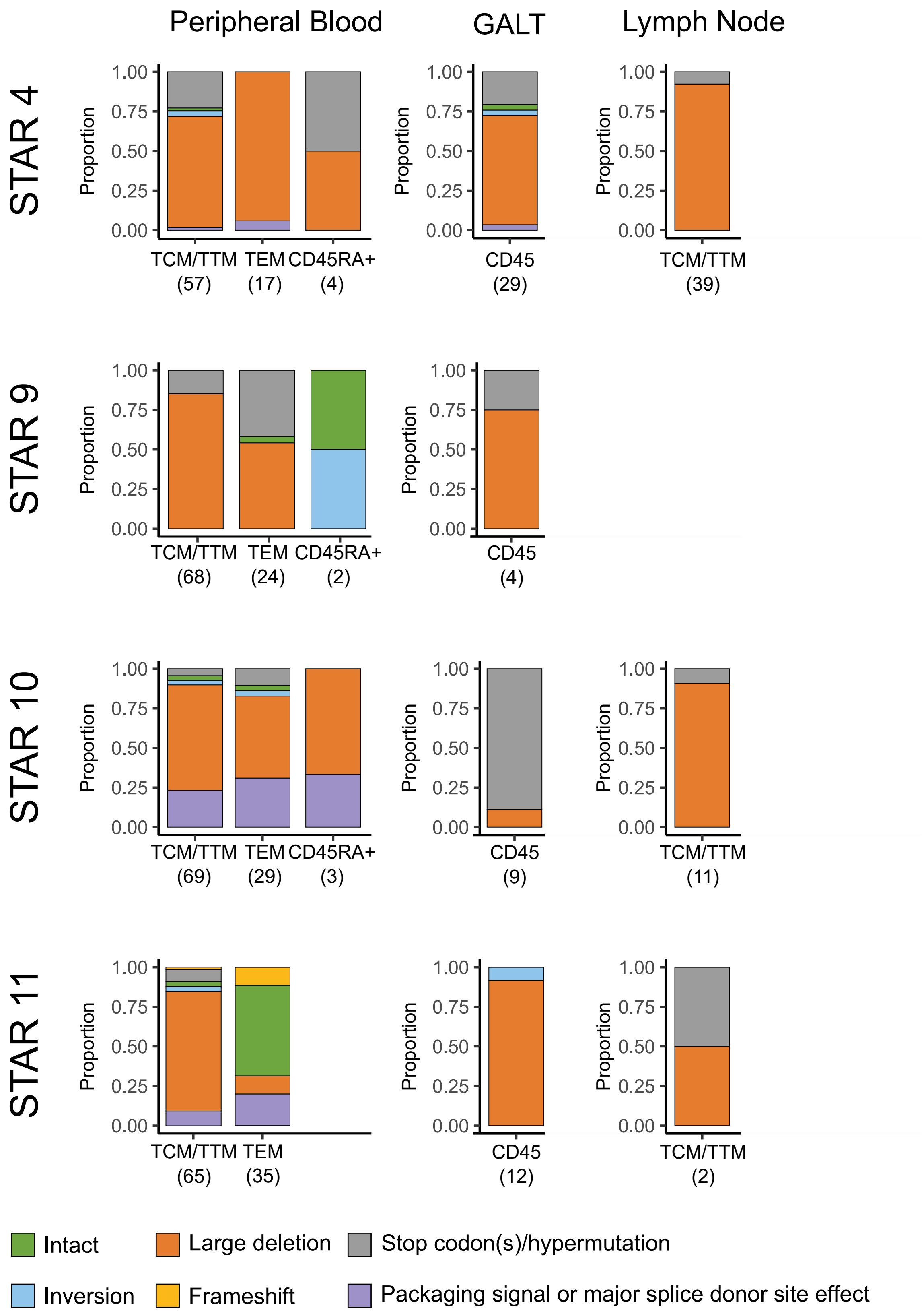


**Supplemental Figure 1:** Proportions of viral categories for each cell subset per anatomical compartment for each participant. The total amount of sequences within each category is shown below in brackets.
TCM/TTM = central/ transitional memory CD4 T cell, TEM = effector memory CD4 T cell, GALT = gut-associated lymphoid tissue.

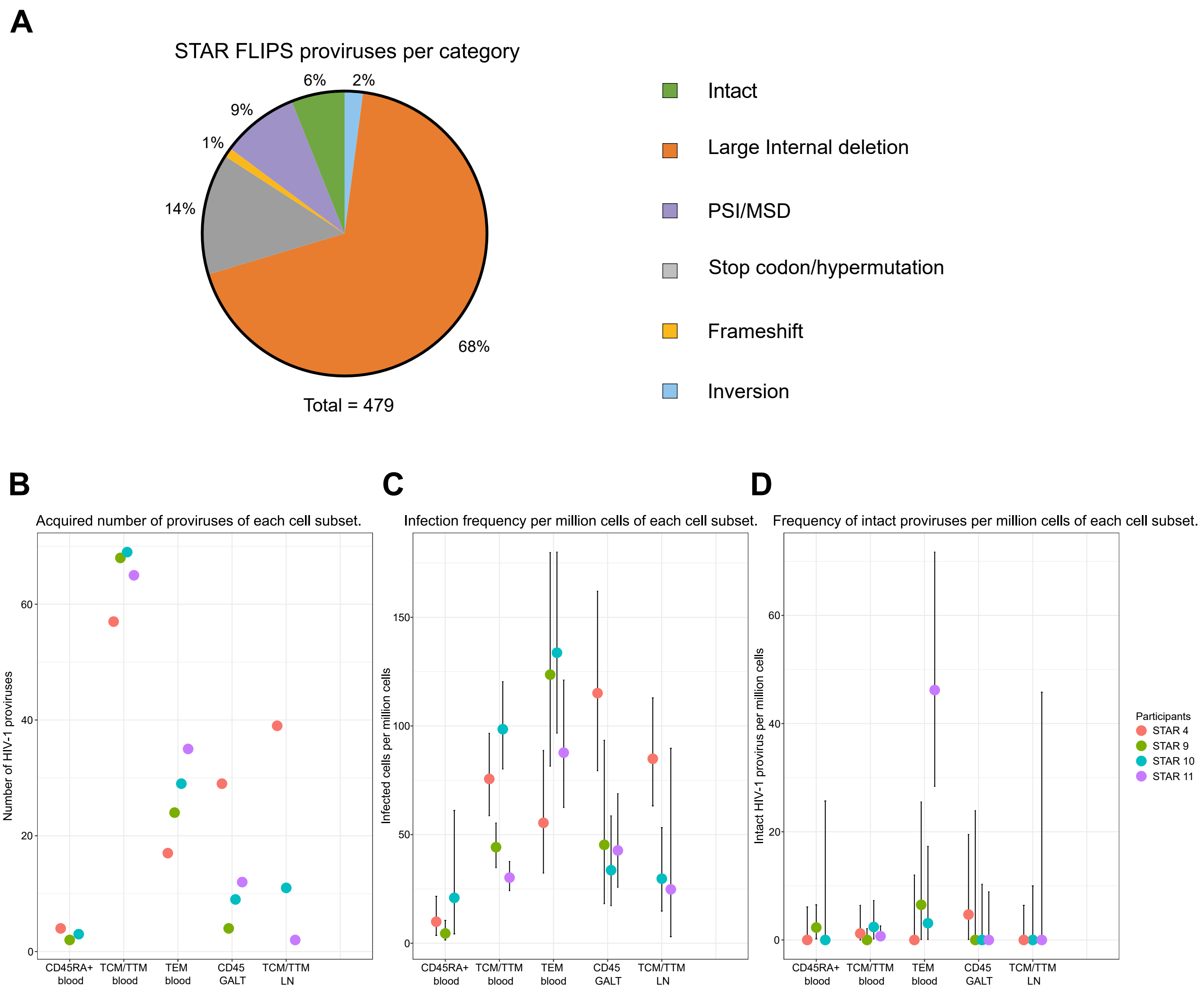


**Supplemental Figure 2:** (A) General overview of the structural categories including all NFL sequences from all participants (B-D) Scatter dot plot representing the infection frequencies for total and intact HIV-1 proviruses shown by cell subset. The legend indicates the color used for each participant. (B) The total number of sequences for each subset per participant. (C) The estimated infection frequencies per million cells with 95% confidence intervals. (D) The estimated infection frequencies of intact proviruses per million cells with 95% confidence intervals. TCM/TTM = central/ transitional memory CD4 T cell, TEM = effector memory CD4 T cell, LN = lymph node, GALT = gut-associated lymphoid tissue.

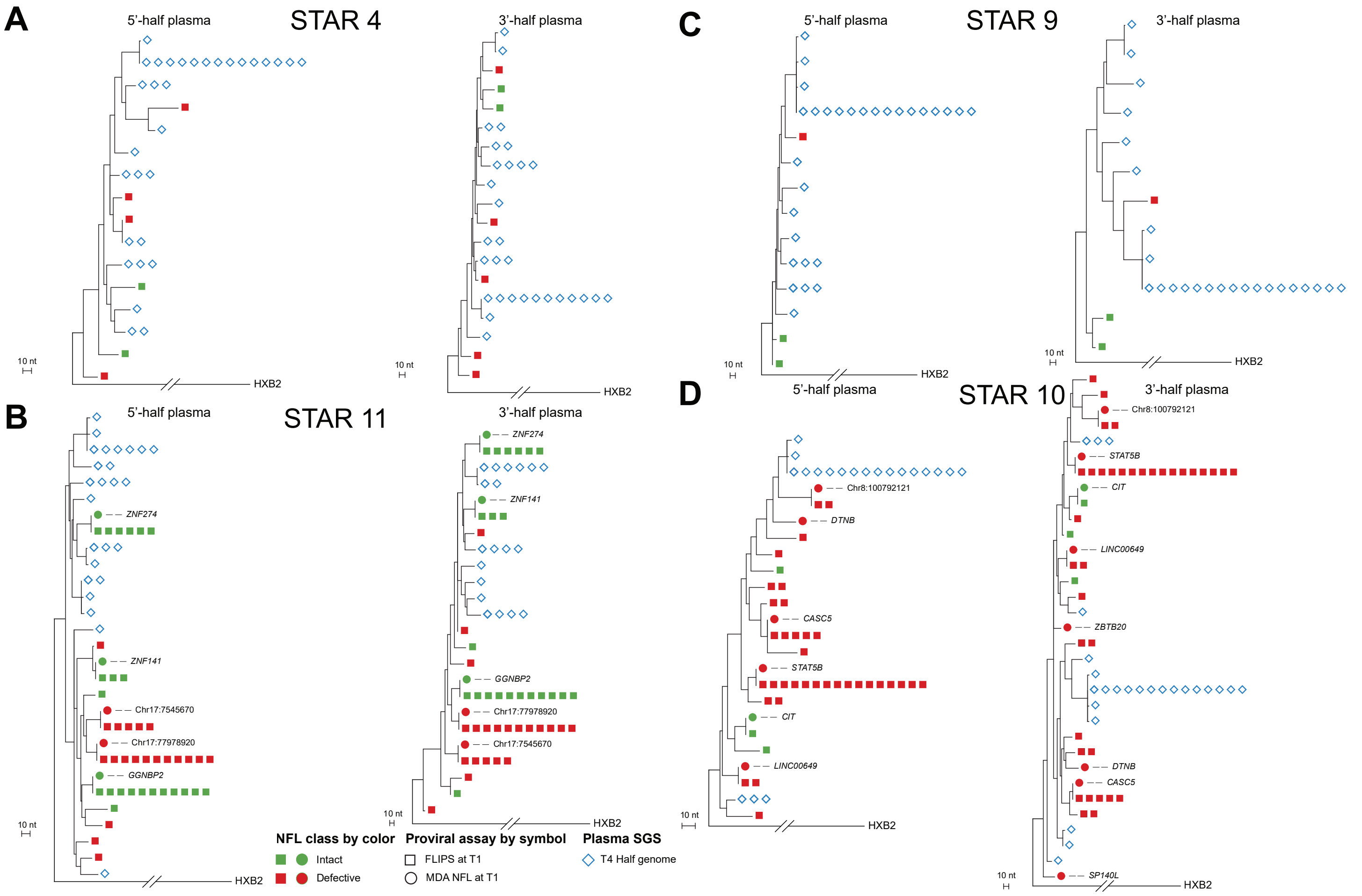


**Supplemental Figure 3:** (A-D) Maximum-likelihood phylogenetic trees based on alignments containing either 5'- or 3'-half rebound (T4) plasma sequences and proviral sequences derived from Full-Length Individual Provirus sequencing (**FLIPS**) and multiple displacement amplification (**MDA**) before analytical treatment interruption (**ATI**, T1). Proviral sequences derived from FLIPS and MDA are shown as squares and circles respectively. The integration sites (**IS**) associated with MDA-derived proviruses are noted if available. Plasma sequences are shown as diamonds. All trees are rooted to the HXB2 reference sequence.

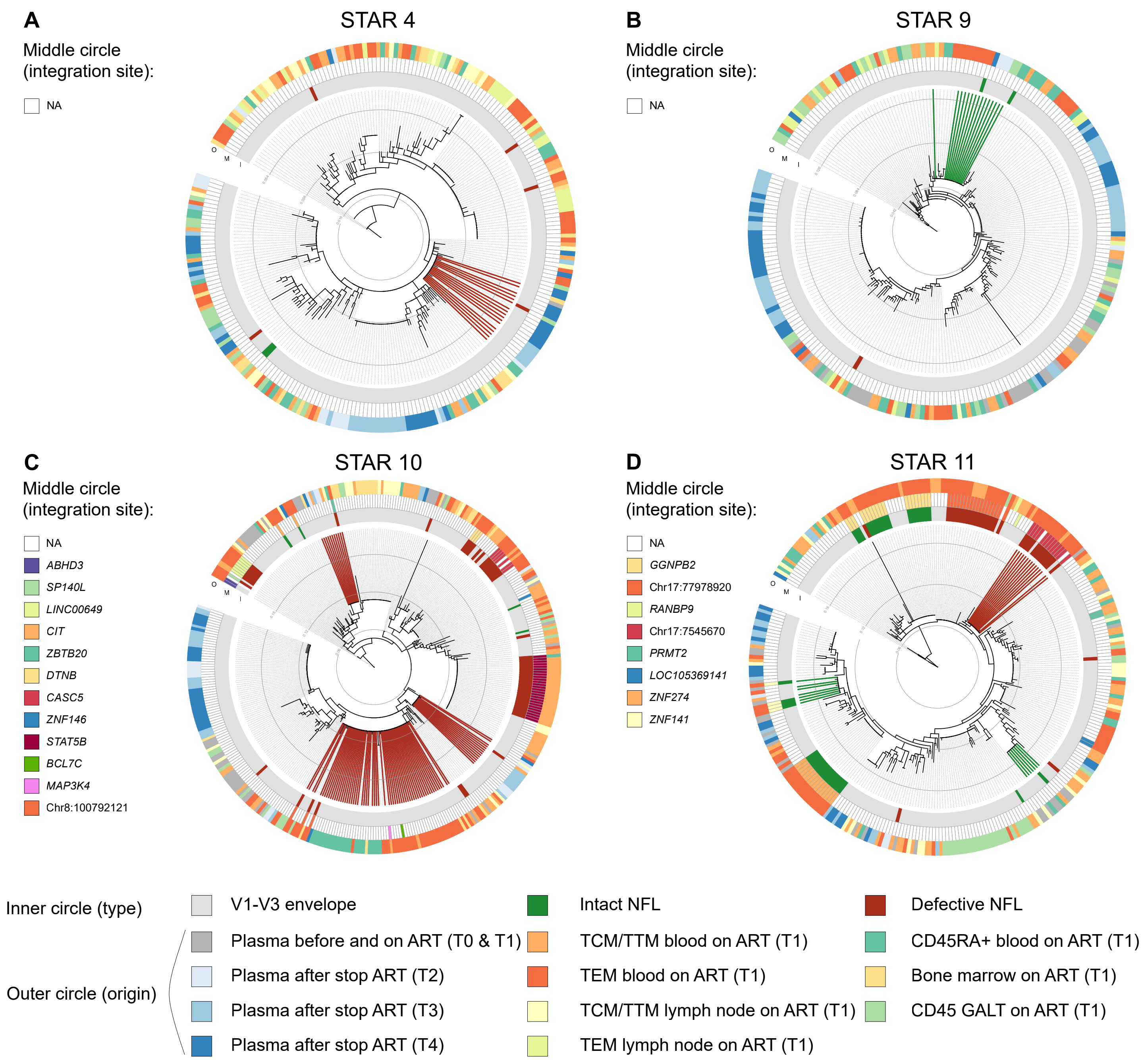


**Supplemental Figure 4 :** Integration sites linked to circular HIV-1 V1-V3 *env* maximum likelihood phylogenetic trees for each participant using all generated proviral and plasma V1-V3 *env* sequences before and during different stages of the ATI shown by cell type origin. The plasma and proviral sequences were obtained either prior ART initiation (timepoint 0, T0), during ART (timepoint 1, T1), 8 to 14 days after analytical treatment interruption (**ATI**) (timepoint 2, T2), at the first detectable viral load (timepoint 3, T3), and at rebound (timepoint 4, T4). The inner circle represents the sequence type, either obtained through single-genome sequencing (**SGS**) of the V1-V3 *env* region shown in grey and V1-V3 *env* trimmed near full-length (**NFL**) genomes in colors indicating their intactness category. The middle circle shows the integration site associated with multiple displacement amplification (**MDA**) derived proviruses if available. The integration sites in the legend are shown in order of appearance on the circle. The outer circle displays the origin (sampling timepoint and/or cell type) of each plasma and proviral sequence. Matches (placed on same branch) of identical V1-V3 *env* regions between plasma and proviral NFL sequences are shown in bold lines, where the line color reflects the intactness category of the matching NFL virus. NA = not available, GALT = gut-associated lymphoid tissue.

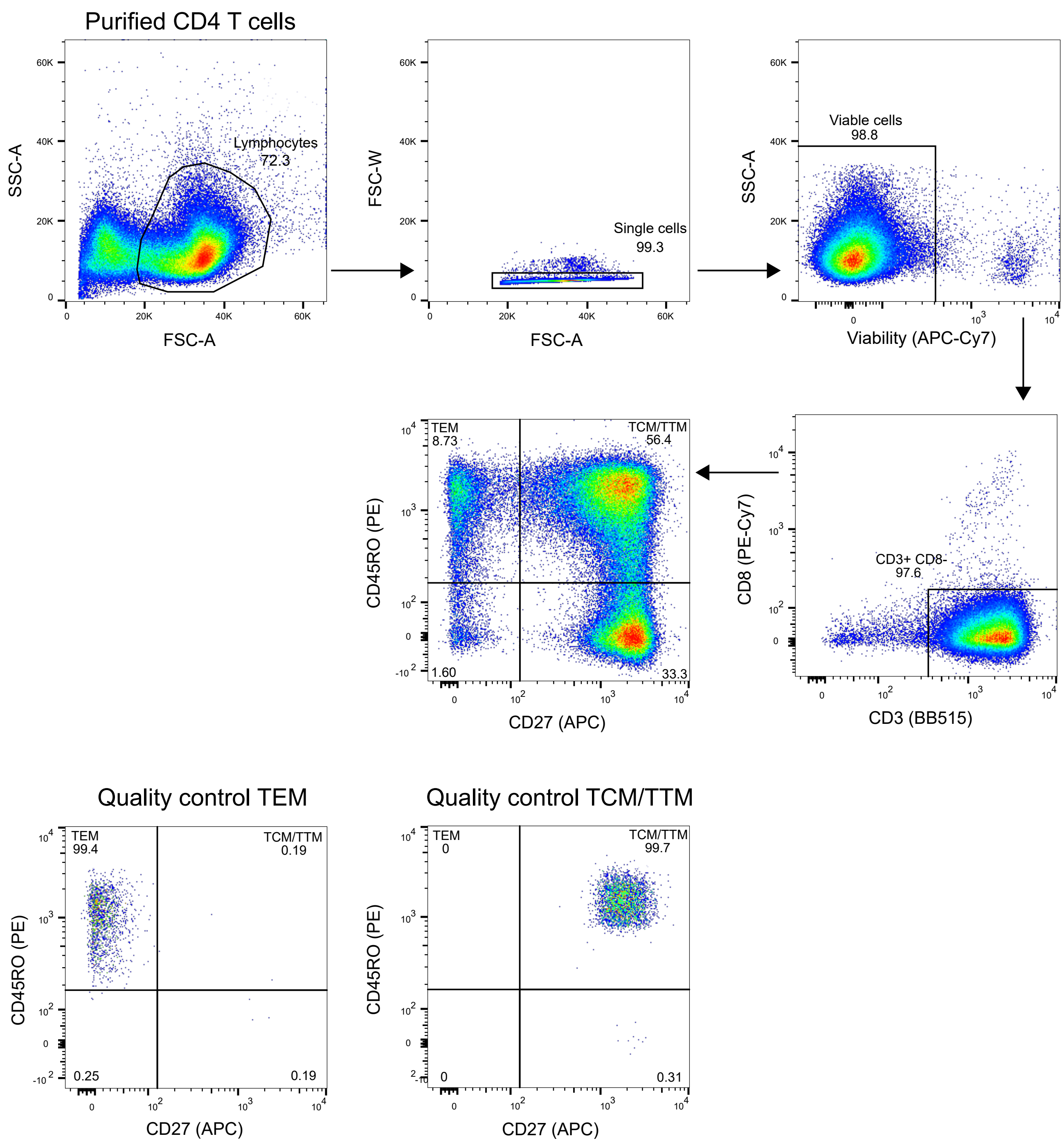


**Supplemental Figure 5:** Gating strategy used for subset sorting from purified CD4 T cells using the CD27 and CD45RO markers. Arrows indicate the sequential gating order. FSC = forward scatter, SSC = side scatter, TCM/TTM = central/ transitional memory CD4 T cell, TEM = effector memory CD4 T cell.

**Supplemental Table 1: Sampled tissues and subsets per assay per participant.** SGS = single-genome sequencing, FLIPS = full-length proviral sequencing, ISLA = integration site loop amplification, MDA = multiple displacement amplification. TCM/TTM = central/transitional memory CD4 T cell, TEM = effector memory CD4 T cell, GALT = gut-associated lymphoid tissue.

| | **Peripheral blood** | | | | | | | | | | | | **GALT** | | | | **Lymph Node** | | | |
|---|---|---|---|---|---|---|---|---|---|---|---|---|---|---|---|---|---|---|---|---|
| | CD45RA+ | | | | TCM/TTM | | | | TEM | | | | CD45 | | | | TCM/TTM | | | |
| | SGS | FLIPS | ISLA | MDA | SGS | FLIPS | ISLA | MDA | SGS | FLIPS | ISLA | MDA | SGS | FLIPS | ISLA | MDA | SGS | FLIPS | ISLA | MDA |
| STAR 4 | x | x | - | - | x | x | - | - | x | x | - | - | x | x | - | - | x | x | - | - |
| STAR 9 | x | x | - | - | x | x | x | x | x | x | x | x | x | x | - | - | x | - | - | - |
| STAR 10 | x | x | - | - | x | x | x | x | x | x | x | x | x | x | - | - | x | x | - | - |
| STAR 11 | x | - | - | - | x | x | x | x | x | x | x | x | x | x | - | - | x | x | - | - |

**Supplemental Table 2: List of integration sites for three study participants: STAR 9, STAR 10 and STAR 11. Integration sites circled in bold were found both in the TCM/TTM and the TEM fraction.** TCM/TTM = central/ transitional memory CD4 T cell, TEM = effector memory CD4 T cell.

*See excel spreadsheet.*

**Supplemental Table 3: Summary of PCRs performed on potentially clonal proviruses.** EIS = expanded identical sequence, MDA = multiple displacement amplification, ISLA = integration site loop amplification.

| | | | | | | | | | | 5-amplicon NFL | | | | | 4-amplicon NFL | | | |
|---|---|---|---|---|---|---|---|---|---|---|---|---|---|---|---|---|---|---|
| **Participant** | **Subset** | **Gene** | **Chromosome** | **Location** | **Orientation WRT gene** | **bulk ISLA count** | **MDA ISLA count** | **V1-V3** | **FLIPS** | **A1mod2** | **A2** | **B2** | **C** | **pol** | **Frag 1** | **Frag 2** | **Frag 3** | **Frag 4** |
| STAR9 | TCM/TTM | *PDCD1LG2* | 9 | 5546773 | Opposite | 0 | 2 | - | - | - | - | - | - | - | NA | NA | NA | NA |
| STAR9 | TCM/TTM | *PAPD5* | 16 | 50160870 | Same | 2 | 0 | NA | NA | NA | NA | NA | NA | NA | NA | NA | NA | NA |
| STAR9 | TCM/TTM | *FAM214A* | 15 | 52602882 | Opposite | 4 | 0 | NA | NA | NA | NA | NA | NA | NA | NA | NA | NA | NA |
| STAR9 | TEM | *FAM214A* | 15 | 52602882 | Opposite | 0 | 2 | - | - | + | - | - | + | - | NA | NA | NA | NA |
| STAR9 | TEM | *NAA25* | 12 | 112070173 | Opposite | 0 | 2 | - | - | - | - | - | - | - | NA | NA | NA | NA |
| STAR9 | TEM | *PPP2CA* | 5 | 134210510 | Opposite | 4 | 2 | - | - | - | - | + | - | - | NA | NA | NA | NA |
| STAR9 | TEM | - | 11 | 334689 | - | 7 | 8 | - | + | - | - | + | + | - | NA | NA | NA | NA |
| STAR9 | TEM | *SMG1P2* | 16 | 29529127 | Opposite | 1 | 1 | - | - | - | - | + | - | - | NA | NA | NA | NA |
| STAR9 | TEM | *ADAM10* | 15 | 58670980 | Opposite | 1 | 1 | - | - | | | | | | NA | NA | NA | NA |
| STAR10 | TCM/TTM | *STAT5B* | 17 | 42264648 | Opposite | 7 | 7 | + | - | + | + | + | + | + | NA | NA | NA | NA |
| STAR10 | TCM/TTM | *CA10* | 17 | 51654848 | Opposite | 3 | 0 | NA | NA | NA | NA | NA | NA | NA | NA | NA | NA | NA |
| STAR10 | TCM/TTM | *ZBTB20* | 3 | 114333246 | Same | 3 | 0 | NA | NA | NA | NA | NA | NA | NA | NA | NA | NA | NA |
| STAR10 | TCM/TTM | *GUSBP2* | 6 | 26873073 | Same | 2 | 0 | NA | NA | NA | NA | NA | NA | NA | NA | NA | NA | NA |
| STAR10 | TCM/TTM | - | 19 | 4072368 | - | 2 | 0 | NA | NA | NA | NA | NA | NA | NA | NA | NA | NA | NA |
| STAR10 | TEM | *SP140L* | 2 | 230350657 | Opposite | 0 | 1 | + | - | - | + | + | + | - | NA | NA | NA | NA |
| STAR10 | TEM | *CIT* | 12 | 119761801 | Same | 0 | 1 | + | - | + | + | + | + | + | NA | NA | NA | NA |
| STAR10 | TEM | - | 8 | 100792122 | - | 0 | 1 | + | - | + | + | + | + | + | NA | NA | NA | NA |
| STAR10 | TEM | *SIK3* | 11 | 116974111 | Same | 0 | 5 | + | - | - | + | - | - | - | NA | NA | NA | NA |
| STAR10 | TEM | *LINC00649* | 21 | 33936112 | Opposite | 0 | 6 | + | - | + | + | + | + | + | NA | NA | NA | NA |
| STAR10 | TEM | *ZBTB20* | 3 | 114333246 | Same | 0 | 1 | + | - | - | + | + | + | - | + | - | - | + |
| STAR10 | TEM | *CASC5* | 15 | 40654989 | Opposite | 0 | 2 | + | - | + | + | + | + | + | NA | NA | NA | NA |
| STAR10 | TEM | *ABHD3* | 18 | 21692430 | Same | 0 | 2 | + | - | - | + | - | - | - | NA | NA | NA | NA |
| STAR10 | TEM | *GATAD2B* | 1 | 153915272 | Same | 0 | 4 | - | - | - | - | - | - | + | NA | NA | NA | NA |
| STAR10 | TEM | *DTNB* | 2 | 25581436 | Opposite | 0 | 3 | + | - | + | + | + | + | + | NA | NA | NA | NA |
| STAR10 | TEM | *VAMP7* | X | 155934835 | Opposite | 0 | 3 | - | - | - | - | - | - | - | NA | NA | NA | NA |
| STAR10 | TEM | *IL6R* | 1 | 154439203 | Same | 0 | 2 | - | - | - | - | - | - | - | NA | NA | NA | NA |
| STAR10 | TEM | *BCL7C* | 16 | 30872736 | Opposite | 0 | 2 | + | - | - | - | - | - | + | NA | NA | NA | NA |
| STAR11 | TCM/TTM | *PRMT2* | 21 | 46639313 | Opposite | 0 | 2 | + | - | + | + | + | + | - | NA | NA | NA | NA |
| STAR11 | TCM/TTM | *UBE2G1* | 17 | 4277762 | Opposite | 0 | 2 | + | - | + | + | + | + | - | NA | NA | NA | NA |
| STAR11 | TEM | *ZNF141* | 4 | 370499 | Opposite | 0 | 1 | + | - | + | + | + | + | + | NA | NA | NA | NA |
| STAR11 | TEM | *ZFC3H1* | 12 | 71648492 | Opposite | 0 | 10 | - | - | - | - | - | - | - | NA | NA | NA | NA |
| STAR11 | TEM | *GGNBP2* | 17 | 36573666 | Same | 0 | 3 | + | - | + | + | + | + | + | NA | NA | NA | NA |
| STAR11 | TEM | - | 17 | 77978920 | - | 0 | 1 | + | - | + | + | + | + | + | NA | NA | NA | NA |
| STAR11 | TEM | - | 17 | 7545670 | - | 0 | 2 | + | - | + | + | + | + | + | NA | NA | NA | NA |
| STAR11 | TEM | - | 6 | 73521230 | - | 0 | 2 | - | - | + | - | - | - | - | NA | NA | NA | NA |
| STAR11 | TEM | *PSMA3-AS1* | 14 | 58283668 | Opposite | 0 | 3 | - | - | - | - | - | - | - | NA | NA | NA | NA |
| STAR11 | TEM | *ZNF274* | 19 | 58194366 | Opposite | 0 | 8 | + | - | + | + | + | + | + | - | - | - | - |
| STAR11 | TEM | *NLRC5* | 16 | 57012433 | Opposite | 0 | 2 | + | - | - | + | + | - | - | NA | NA | NA | NA |
| STAR11 | TEM | *EHMT1* | 9 | 137713251 | Opposite | 0 | 2 | - | - | - | - | - | - | + | NA | NA | NA | NA |

**Supplemental Table 4: Number of intact and defective sequences retrieved from analyzed cells for each cell subset and anatomical compartments from all participants.** TCM/TTM = central/transitional memory CD4 T cell, TEM = effector memory CD4 T cell.

| | | | Participant | | | |
|---|---|---|---|---|---|---|
| | | | STAR 4 | STAR 9 | STAR 10 | STAR 11 |
| Peripheral blood | CD45RA+ | Defective | 4 | 1 | 3 | - |
| | | Intact | 0 | 1 | 0 | - |
| | | No of cells analysed | 604,263 | 1,109,706 | 143,400 | - |
| | TCM/TTM | Defective | 56 | 68 | 67 | 63 |
| | | Intact | 1 | 0 | 2 | 2 |
| | | No of cells analysed | 869,109 | 1,719,541 | 992,146 | 2,886,308 |
| | TEM | Defective | 17 | 23 | 28 | 15 |
| | | Intact | 0 | 1 | 1 | 20 |
| | | No of cells analysed | 306,771 | 218,420 | 321,708 | 430,579 |
| GALT | CD45 | Defective | 28 | 4 | 9 | 12 |
| | | Intact | 1 | 0 | 0 | 0 |
| | | No of cells analysed | 286,062 | 154,391 | 357,444 | 413,481 |
| Lymph node | TCM/TTM | Defective | 39 | - | 11 | 2 |
| | | Intact | 0 | - | 0 | 0 |
| | | No of cells analysed | 573,680 | - | 369,893 | 80,525 |

**Supplemental Table 5: Clonal prediction scores and nucleotide distances of each participant.** CPS = Clonal Prediction Score, NFL = near full-length, FLIPS = full-length proviral sequencing.

| Participant | Total NFL FLIPS proviruses | Total unique NFL FLIPS proviruses | Total hypothetical V1-V3 detectable proviruses | Total unique hypothetical V1-V3 detectable proviruses | CPS (%) | Nucleotide diversity of V1-V3 region |
|---|---|---|---|---|---|---|
| STAR 4 | 146 | 114 | 27 | 26 | 96% | 0.0101 |
| STAR 9 | 98 | 71 | 14 | 14 | 100% | 0.01724 |
| STAR 10 | 121 | 60 | 22 | 21 | 95% | 0.01428 |
| STAR 11 | 114 | 77 | 17 | 17 | 100% | 0.01641 |

### Supplemental Table 6: Participant characteristics.

| Participant | Gender | Age (y) | CD4 nadir (cells/µl) | VL zenith (log10 copies/mL) | CD4/CD8 at T1 | CD4 at T1 (cells/µl) | Time since primo-infection (y) | Time on cART before ATI (y) | Time to viral rebound (d) | VL at T0 (copies/mL) | VL at T0 (copies/mL) | VL at T1 (copies/mL) | VL at T2 (copies/mL) | VL at T3 (copies/mL) | VL at T4 (copies/mL) | HIV-1 subtype | Viral tropism |
|---|---|---|---|---|---|---|---|---|---|---|---|---|---|---|---|---|---|
| STAR 4 | M | 36 | 142 | 4.31 | 1.06 | 756 | NA | 11 | 20 | NA | NA | <20 | <20 | 790 | 36500 | B | R5 |
| STAR 9 | M | 32 | 405 | 5.01 | 0.84 | 918 | 9 | 6 | 21 | NA | 30115 | <20 | <20 | 101 | 1967 | B | R5 |
| STAR 10 | M | 54 | 327 | 5.49 | 0.66 | 736 | 29 | 11 | 21 | 82110 | 3190 | <20 | <20 | 683 | 4450 | B | R5 |
| STAR 11 | M | 37 | 432 | 3.62 | 0.94 | 911 | 9 | 2 | 28 | 4160 | 12589 | <20 | <20 | 202 | 1790 | B | R5 |

#### Supplemental Table 7: Primers used for 5' ISLA.

| Step | Primer name | Sequence (5' to 3') |
|---|---|---|
| Linear extension | UTR.629.R | CCCTGTTCGGGCGCCACTGCTA |
| Decamer binding and extension | decaU3R.3 | GTTCTGCCAATCAGGGAAGTAGCCTTGTGTGTNNNNNNNNNN |
| PCR 1 | U3R.1 | GGCTCAACTGGTACTAGCTTGAAGCACCATCCAAAG |
| PCR 2 | U3R.2 | GGATATCTGATCCCTGGCCCTGGTGTGTAGTT |
| PCR 3 | U3R.3 | GTTCTGCCAATCAGGGAAGTAGCCTTGTGTGT |
| Sanger sequencing | U3R.4 | CCCACAGATCAAGGATATCTTGTCT |
